# Caveolae set levels of epithelial monolayer tension to eliminate tumor cells

**DOI:** 10.1101/632802

**Authors:** Jessica L. Teo, Guillermo A. Gomez, Ivar Noordstra, Suzie Verma, Vanesa Tomatis, Bipul R. Acharya, Lakshmi Balasubramaniam, Hiroko Katsuno-Kambe, Rachel Templin, Kerrie-Ann McMahon, Robert J. Ju, Samantha J. Stebhens, Benoit Ladoux, Robert G. Parton, Alpha S. Yap

**Affiliations:** Division of Cell Biology and Molecular Medicine, Institute for Molecular Bioscience, The University of Queensland, St. Lucia, Brisbane, Queensland, Australia 4072; Centre for Cancer Biology, SA Pathology and University of South Australia, Adelaide SA 5000.; Institut Jacques Monod, CNRS, UMR 7592, Université de Paris, 75013 Paris, France; The University of Queensland, The University of Queensland Diamantina Institute, Translational Research Institute, Woolloongabba, Brisbane, QLD 4102, Australia; Centre for Microscopy and Microanalysis, The University of Queensland, St. Lucia, Brisbane, Queensland, Australia 4072

**Keywords:** Caveolae, Ras, Cell extrusion, phosphoinostides, actomyosin

## Abstract

Mechanical tension governs epithelial morphogenesis and homeostasis, but its regulation remains poorly understood. Tension is commonly contractile, arising when the actomyosin cortices of cells are mechanically coupled together by cadherin adhesion. Here we report that caveolae control levels of epithelial tension and show that this is necessary for oncogene-transfected cells to be eliminated by apical extrusion. Depletion of caveolin-1 (CAV1) in the surrounding epithelium, but not in the oncogene-expressing cells, blocked extrusion leading to the retention and proliferation of transformed cells within the monolayer. Tensile stress was aberrantly elevated in CAV1-depleted monolayers due to elevated levels of phosphoinositide-4,5-bisphosphate (PtdIns(4,5)*P*_2_) causing increased recruitment of the formin, FMNL2. Oncogenic extrusion was restored to CAV1-deficient monolayers when tension was corrected by depleting FMNL2, blocking PtdIns(4,5)*P*_2_, or disabling the interaction between FMNL2 and PtdIns(4,5)*P*_2_. Thus, by controlling lipid signalling to the actin cytoskeleton, caveolae regulate mechanical tension for epithelial homeostasis.

## Introduction

Epithelia commonly display mechanical tension (Fernandez-Gonzalez et al., 2009; Martin et al., 2010; Ratheesh et al., 2012). An important source of this tension arises when the contractile actomyosin cortices of individual cells are mechanically coupled together by cadherin cell-cell adhesion (Acharya et al., 2018; Charras and Yap, 2018; le Duc et al., 2010). Cadherin signalling also modulates the activity of the actomyosin apparatus itself (Kovacs et al., 2002; Wu et al., 2014). Together, these processes create a network of tensile adherens junctions that extend throughout epithelial sheets. During development, this tissue tension exerts morphogenetic effects that are apparent on different length scales. Local patterns of tension participate in local cellular rearrangements, such as neighbour exchange and cellular ingression (Fernandez-Gonzalez et al., 2009; Kobb et al., 2017; Rauzi et al., 2008; Samarage et al., 2015); but tissue-scale anisotropies have also been demonstrated, which drive large-scale patterns of folding in the embryo (Martin et al., 2010; Rauzi et al., 2015). These instabilities in tension are controlled by the spatially and temporally restricted expression of developmental signals, such as the transcription factors Twist and Snail (Martin et al., 2010), whose impact is ultimately expressed through cytoskeletal regulators, including the RhoA GTPase.

Junctional and tissue tension is also evident in morphologically stable epithelia, such as confluent cultured monolayers (Acharya et al., 2018; Trepat and Sahai, 2018; Trepat et al., 2009; Wu et al., 2014). Here, tension participates in physiological and homeostatic processes, such as barrier permeability (Turner et al., 1997) and the elimination of cells by apical extrusion (Saw et al., 2017). Interestingly, the level of junctional tension appears to be functionally important. For example, epithelial permeability is compromised when tension is reduced (Acharya et al., 2017) and also when it is increased (Fanning et al., 2012; Tsukita et al., 2009). This implies that there must be cellular mechanisms that determine (or “set”) the steady-state level of junctional tension necessary for epithelial homeostasis. What those mechanisms may be is not yet known.

Caveolae are plasma membrane organelles that have recently been implicated in cellular mechanics. These flask-shaped membrane invaginations are formed through the action of the integral membrane proteins, caveolins, and their coat proteins, cavins (Parton and Collins, 2016; Parton and del Pozo, 2013). Caveolae flatten and disassemble when mechanical stress is applied to cells; this limits the degree to which membrane tension is increased by mechanical stress (Sinha et al., 2011). Thus, caveolae are mechanosensitive and responsible for eliciting compensatory mechanical responses. Their ability to buffer membrane tension has been attributed to the release of a membrane reservoir when caveolae disassemble (Sinha et al., 2011). Caveolae also have other diverse effects which can potentially influence cell signalling and mechanics, including control of the lipid composition and organization of the plasma membrane (Ariotti et al., 2014). However, whether these or other potential pathways are utilized to regulate tissue tension – and what role they may play in epithelial homeostasis - has not been tested.

We now report that caveolae set levels of monolayer tension by controlling junctional contractility. This affects the ability of epithelia to eliminate oncogene-expressing cells by apical cell extrusion. The latter process (also called oncogenic cell extrusion) occurs when minorities of oncogene-expressing cells are surrounded by oncogene-free neighbours (Hogan et al., 2009; Kajita et al., 2014; Leung and Brugge, 2012). It is compromised when the mechanical integrity of the surrounding epithelium is disrupted by E-cadherin depletion (Hogan et al., 2009) and when apical-basal patterns of mechanical tension are altered at adherens junctions (Wu et al., 2014). This homeostatic response of extrusion is elicited by many different oncogenes and in a range of epithelia, suggesting that it is a common epithelial response to early events of transformation. We show that oncogenic extrusion is compromised in CAV1-deficient epithelia. Strikingly, caveolae are selectively required in the neighbouring cells, rather than in oncogene-expressing cells. This is due to an abnormal elevation of junctional tension in the epithelium that arises from aberrant actin regulation by phosphoinositide-4,5-bisphosphate (PtdIns(4,5)*P*_2_).

## RESULTS

### Caveolin-1 supports oncogenic extrusion

CAV1 is a fundamental structural element of caveolae that is necessary for their biogenesis (Hayer et al., 2010a; Parton and del Pozo, 2013). Given the proposed role of caveolae in regulating the mechanical properties of cells (Sinha et al., 2011) and the role of tissue mechanics in apical extrusion (Wu et al., 2014, we tested how reducing CAV1 levels affected the extrusion of oncogene-expressing cells in AML12 liver epithelial monolayers. CAV1 KD depleted its protein expression by >90% (Fig S1a) and reduced the number of caveolae that were detectable, as shown by immunofluorescence microscopy for the caveolar coat protein, cavin-1 (Fig S1e). Deconvolution confocal microscopy further revealed that CAV1 co-accumulated with E-cadherin at cell-cell junctions in wild-type AML12 cells (Fig S1c), suggesting that caveolae might preferentially localize to adherens junctions (AJ). This was confirmed both by thin section electron microscopy (Fig S1b) and focused ion beam scanning electron microscopy (Fig S1d), which showed clear concentration of caveolae close to the AJ at the apical region of the lateral border. Interestingly, caveolae clustered at AJ to form prominent rosettes (Fig S1d), a mode of organization that is particularly sensitive to membrane tension (Golani et al., 2019).

To induce oncogenic extrusion, H-Ras^V12^ was expressed in groups of 1-2 cells that were surrounded by untransfected cells (Fig 1a). Under these mosaic conditions, H-Ras^V12^ cells were apically extruded from the monolayers at ~ 5-times the rate seen when a control mCherry transgene was expressed (Fig 1b). In contrast, apical extrusion of H-Ras^V12^ cells was substantially reduced by CAV1 KD (Fig 1a,b). The specificity of the RNAi was confirmed by reconstitution with an RNAi-resistant CAV1 transgene (Fig 1a,b). CAV1 KD also inhibited the apical extrusion of cells expressing oncogenic c-Src^Y527F^ (Fig 1c) or K-Ras^G12V^ (Fig S1g). Thus, CAV1 was necessary for the extrusion of cells expressing a wide variety of oncogenes, suggesting that it might generally influence oncogenic extrusion.

**Figure 1.**
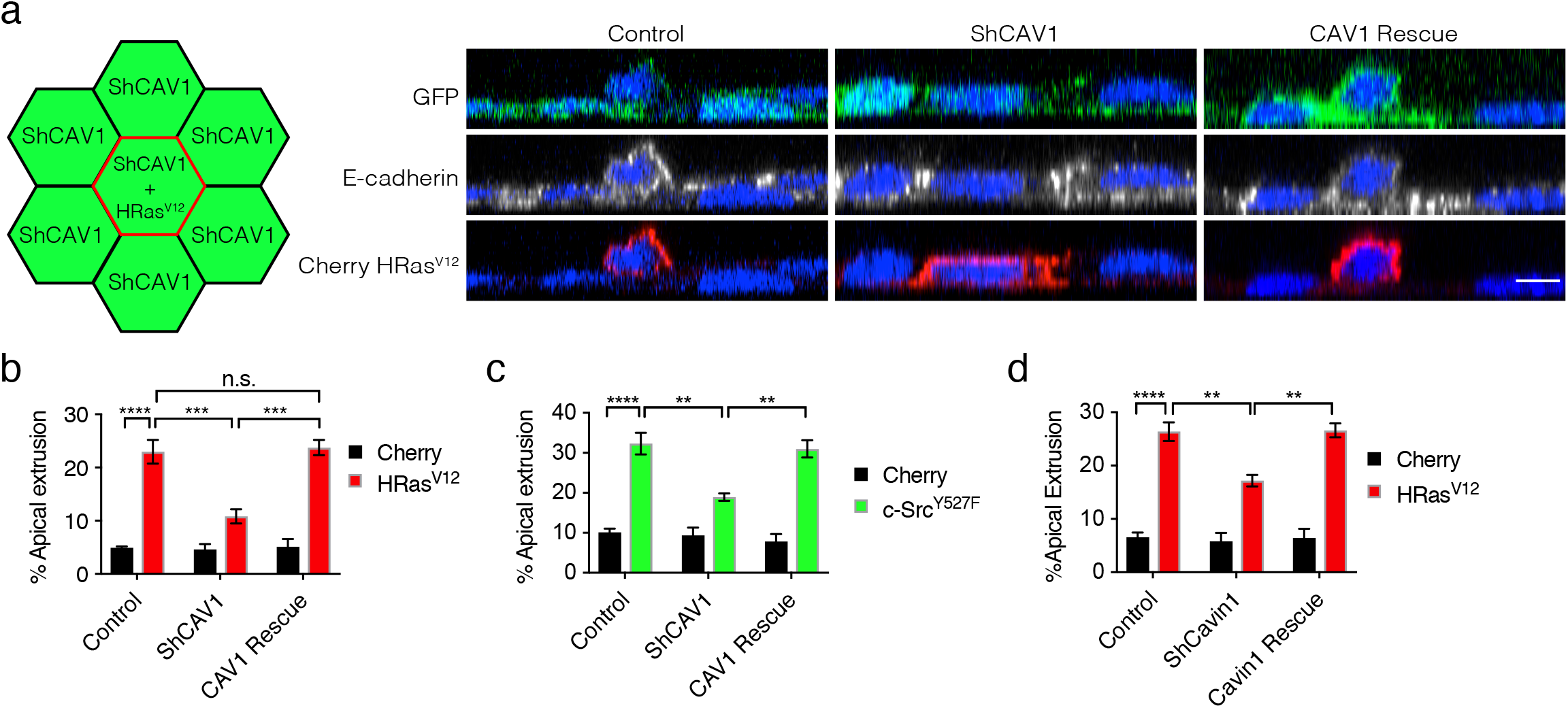
Caveolin-1 supports oncogenic cell extrusion. (a) Schematic illustration and representative XZ images of single HRas^V12^-expressing cells within GFP control, ShCAV1 GFP and CAV1 Rescue GFP epithelial monolayers. (b,c) Quantification of apical cell extrusion of single cells expressing (b) HRas^V12^ or (C) c-Src^Y527F^ within GFP control, ShCAV1 GFP and CAV1 Rescue GFP epithelial monolayers. Soluble mCherry was expressed as a control for the mCherry-tagged oncogenes. (d) Role for Cavin 1 in oncogenic cell extrusion. Quantification of apical cell extrusion of single cells expressing HRas^V12^ within GFP control, ShCavin1 GFP and Cavin1 Rescue GFP epithelial monolayers. All data are means ± s.e.m; n.s, not significant; \**P*<0.05, \*\**P*<0.01, \*\*\**P*<0.001, \*\*\*\**P*<0.0001; calculated from N=3 independent experiments analysed with two-way ANOVA Tukey’s multiple comparisons test. Scale bar, 10µm.

Although CAV1 KD reduced caveolae in our cells (Fig S1a, e), CAV1 can also have caveola-independent effects (Hayer et al., 2010b; Hill et al., 2008). Therefore, we depleted Cavin1 (Fig S1f), which is also necessary for caveolar integrity (Hill et al., 2008; Liu et al., 2008). Cavin1 shRNA reduced the apical extrusion of H-Ras^V12^ (Fig 1d) and c-Src^Y527F^-expressing cells (Fig S1h) as effectively as did CAV1 KD. As CAV1 levels were not altered by Cavin1 KD under these conditions (Fig S1i), this argues that oncogenic extrusion requires intact caveolae, not just the presence of CAV1.

### Oncogenic extrusion specifically requires caveolae in the surrounding epithelium

Apical extrusion arises from the interaction between oncogene-expressing cells and the non-transformed cells of the surrounding epithelium (Hogan et al., 2009; Kajita et al., 2014). To understand which of these cells required caveolae for effective extrusion we employed mixing experiments that depleted CAV1 in selected populations (Fig 2). First, we depleted CAV1 in the oncogene-expressing cells. For this, we expressed H-Ras^V12^ in CAV1 KD cells, then mixed them with wild-type (WT) cells at a ratio (1:20) where the majority of H-Ras^V12^/CAV1^KD^ cells were surrounded by WT cells (Fig 2a). Surprisingly, H-Ras^V12^/CAV1^KD^ cells underwent extrusion at rates comparable to those seen when H-Ras^V12^ was expressed on a WT background (Fig 2b). Similarly, CAV1 KD in cells expressing c-Src^Y527F^ did not affect their rates of extrusion, so long as the surrounding cell population was wild-type (Fig 2c). Therefore, CAV1 was not required in the oncogene-expressing cell for extrusion to occur.

**Figure 2.**
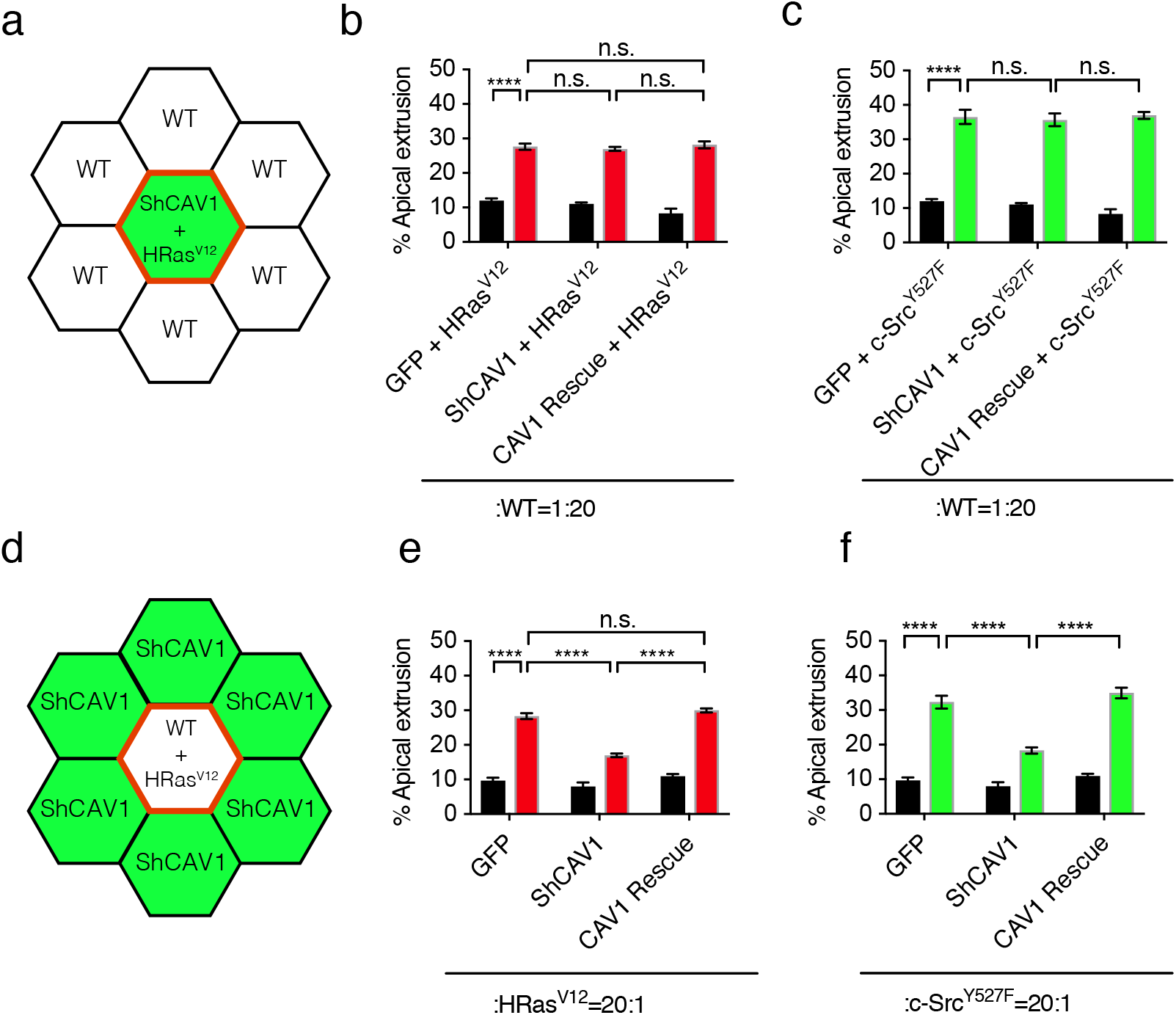
CAV1 is selectively required in the surrounding epithelium for oncogenic cell extrusion. (a-c) Effect of depleting CAV1 selectively in oncogene-expressing cells. (a) Schematic of the experimental design. Oncogenes were expressed in GFP control, ShCAV1 GFP or CAV1 Rescue GFP cells and mixed with a majority of wild-type cells. (b,c) Quantification of apical cell extrusion when CAV1 was manipulated in either (b) HRas^V12^ or (c) c-Src^Y527F^-expressing cells. (d-f) Effect of depleting CAV1 selectively in the surrounding epithelium. (d) Schematic of the mixing design. Oncogenes were expressed on a CAV1 wild-type background and CAV1 manipulated in the surrounding cells. (e,f) Quantification of apical cell extrusion for (e) HRas^V12^ or (f) c-Src^Y527F^. All data are means ± s.e.m; n.s, not significant; \**P*<0.05, \*\**P*<0.01, \*\*\**P*<0.001, \*\*\*\**P*<0.0001; calculated from N=3 independent experiments analysed with two-way ANOVA Tukey’s multiple comparisons test.

This suggested that effective apical extrusion might only require caveolae to be present in the surrounding epithelial population. To test this, we mixed WT cells expressing H-Ras^V12^ (H-Ras^V12^/WT) with CAV1 KD cells forming the surrounding epithelium (Fig 2d). Here we found that extrusion was inhibited (Fig 2e). Similarly, extrusion of c-Src^Y527F^/WT cells was inhibited when their surrounding cells were CAV1 depleted (Fig 2f). Together, these observations indicate that CAV1, and by implication, caveolae, are selectively required in the surrounding epithelium for oncogenic extrusion to occur. This suggested that caveolae may influence some general property of the epithelial monolayer that is necessary to permit oncogenic extrusion to occur.

### CAV1 depletion increases epithelial monolayer tension

Extrusion is influenced by patterns of contractile tension at cell-cell contacts (Wu et al., 2014). We therefore focused on the possibility that CAV1 depletion might generally alter the biomechanics of AML12 monolayers. First, we used Bayesian Inference Stress microscopy (BISM (Nier et al., 2016)) to assess the supracellular or tissue-level stresses in monolayers. For this, traction force microscopy (TFM) was performed on cells grown to confluence on flexible PDMS substrata coated with fluorescent beads, and the cellular-level stresses were back-calculated from the TFM data. Wild-type AML12 cells showed heterogeneity, with regions that were tensile and others compressive. Overall, the average isotropic stress was compressive in these confluent cultures (Fig 3a). Strikingly, however, average isotropic stress became tensile in CAV1 KD monolayers at similar densities and this was restored to resemble wild-type cells by reconstitution of CAV1 (Fig 3a). In addition, treating CAV1 KD cells with the ROCK inhibitor Y27632 restored isotropic stress to control levels (Fig 3a). This suggested that the tensile stress was due to an increase in cellular contractility. Therefore, CAV1 depletion increased tissue-level tension in the monolayers.

**Figure 3.**
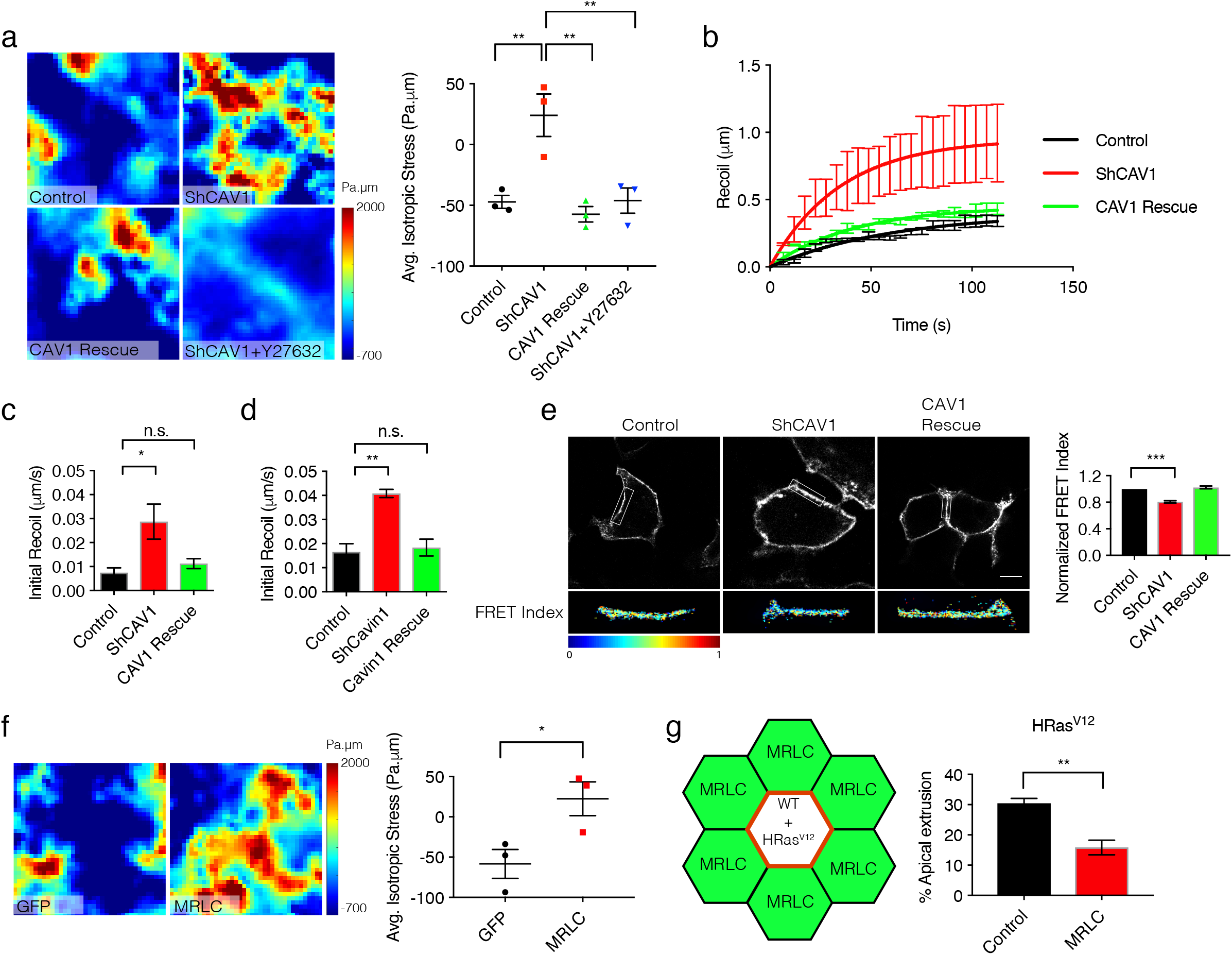
CAV1 depletion increases epithelial monolayer tension. (a) Effect of CAV1 on monolayer stress measured by Bayesian Inversion Stress Microscopy. Representative heat map and quantification of average isotropic stress across 12hrs. (b-d) Junctional tension measured by initial recoil after laser ablation. (b-c) Effect of CAV1 depletion. Representative recoil curves for E-cadherin-mCherry (b) and (c) quantitation of initial recoil velocity. (d) Effect of Cavin1 shRNA on initial recoil velocity. (e)Molecular tension at AJ inferred from a FRET-based α-catenin-tension sensor (TS). Representative images of junctional localization and FRET index from α-catenin-TS; and quantitated normalized FRET Index. (f) Exogenous MRLC-EGFP increases monolayer stress: representative heat map and quantification of average isotropic stress. GFP alone was expressed as a control. (g) Apical oncogenic cell extrusion in mixing experiments when inducible HRas^V12^-expressing WT cells were mixed with a majority of GFP control or MRLC-EGFP-expressing cells. Schematic illustration and quantification of apical extrusion. All data are means ± s.e.m; n.s, not significant; \**P*<0.05, \*\**P*<0.01, \*\*\**P*<0.001, \*\*\*\**P*<0.0001; calculated from N=3 independent experiments analysed with one-way ANOVA Tukey’s multiple comparisons test except (f) and (g) which were analysed by Student’s *t*-test. Scale bar, 10µm.

A major source of tissue tension in epithelia comes from the mechanical coupling of the actomyosin cytoskeleton to cell-cell AJ (Charras and Yap, 2018). We tested if caveolae might regulate junctional tension by measuring recoil after E-cadherin junctions were cut with a laser. As previously reported (Ratheesh et al., 2012), junctions displayed immediate recoil after laser ablation, confirming that they were under tension at steady-state (Fig 3b-d). Recoil velocities were significantly and specifically increased by either CAV1 KD (Fig 3b,c) or cavin 1 KD (Fig 3d, S2a). This increase in recoil was attributable to an increase in junctional tension, junctional friction being unchanged, as inferred from *k*-values when recoil was modelled as a Kelvin-Vogt fiber (Table S1). To corroborate this, we measured molecular level-tension at AJ using an α-catenin transgene that incorporates a FRET-based tension-sensor (TS) module (α-cat-TS) (Acharya et al., 2017). Energy transfer across α-cat-TS was reduced in CAV1 KD cells compared with controls (Fig 3e), indicating an increase in molecular-level tension. Together, these findings reveal that junctional and tissue tension is aberrantly elevated in CAV1 KD cells.

This led us to consider whether increased mechanical tension in the surrounding epithelium might inhibit oncogenic extrusion in CAV1 KD cells. As a first, in-principle test of this hypothesis, we over-expressed the myosin II regulatory light chain (MRLC-EGFP) in wild-type AML12 cells. MRLC-EGFP localized to cell-cell junctions (Fig S2c) and increased total cellular levels of pMLC (Fig S2d, e, reflecting both endogenous MRLC and the transgene), suggesting that it increased the pool of active Myosin II. This was accompanied by increased contractile tension as reflected in junctional recoil (Fig S2f, g) and monolayer stress (Fig 3f). However, caveolae remained detectable when immunostained for cavin-1 (Fig S2h). MRLC-EGFP-AML12 cells were then mixed as neighbour cells with a minority of cells that expressed oncogenes on a WT-background (Fig 3g). In order to refine their temporal expression, we expressed HRas^V12^ or c-Src^Y527F^ under the control of the Tet^ON^ promoter (Fig S2b, i). Oncogene-transfected cells were mixed with MRLC-EGFP-or control cells (1:20), grown to confluence in the suppressed state, before addition of doxycycline to induce expression (2 µg/ml, 24-48 h; Fig S2b, i). Strikingly, extrusion of both Tet^ON^-HRas^V12^ cells (Fig 3g) and Tet^ON^-c-Src^Y527F^ cells (Fig S2j) was significantly reduced when they were surrounded by MRLC-EGFP-AML12 cells, compared with a control epithelium (Fig 3g, S2j). This indicated that increasing tissue tension had the capacity to inhibit oncogenic extrusion even when epithelia were wild-type for CAV1. To test if this could explain what we observed with CAV1 deficiency, it was then necessary to identify how CAV1 depletion increased contractile tension.

### A formin is responsible for increasing tension at CAV1 KD junctions

Contractile tension is driven by the actomyosin cortex (Chugh and Paluch, 2018). However, despite increased junctional tension, CAV1 KD did not change junctional levels of non-muscle myosin II (Fig S3a,b). Nor was the expression of E-cadherin or its junctional dynamics altered by CAV1 KD (Fig S3c-g). Instead, F-actin levels detected by quantitation of phalloidin intensity at junctions were subtly but consistently elevated in CAV1 KD cells (Fig 4a). This suggested that actin regulation might be altered by CAV1 depletion.

**Figure 4.**
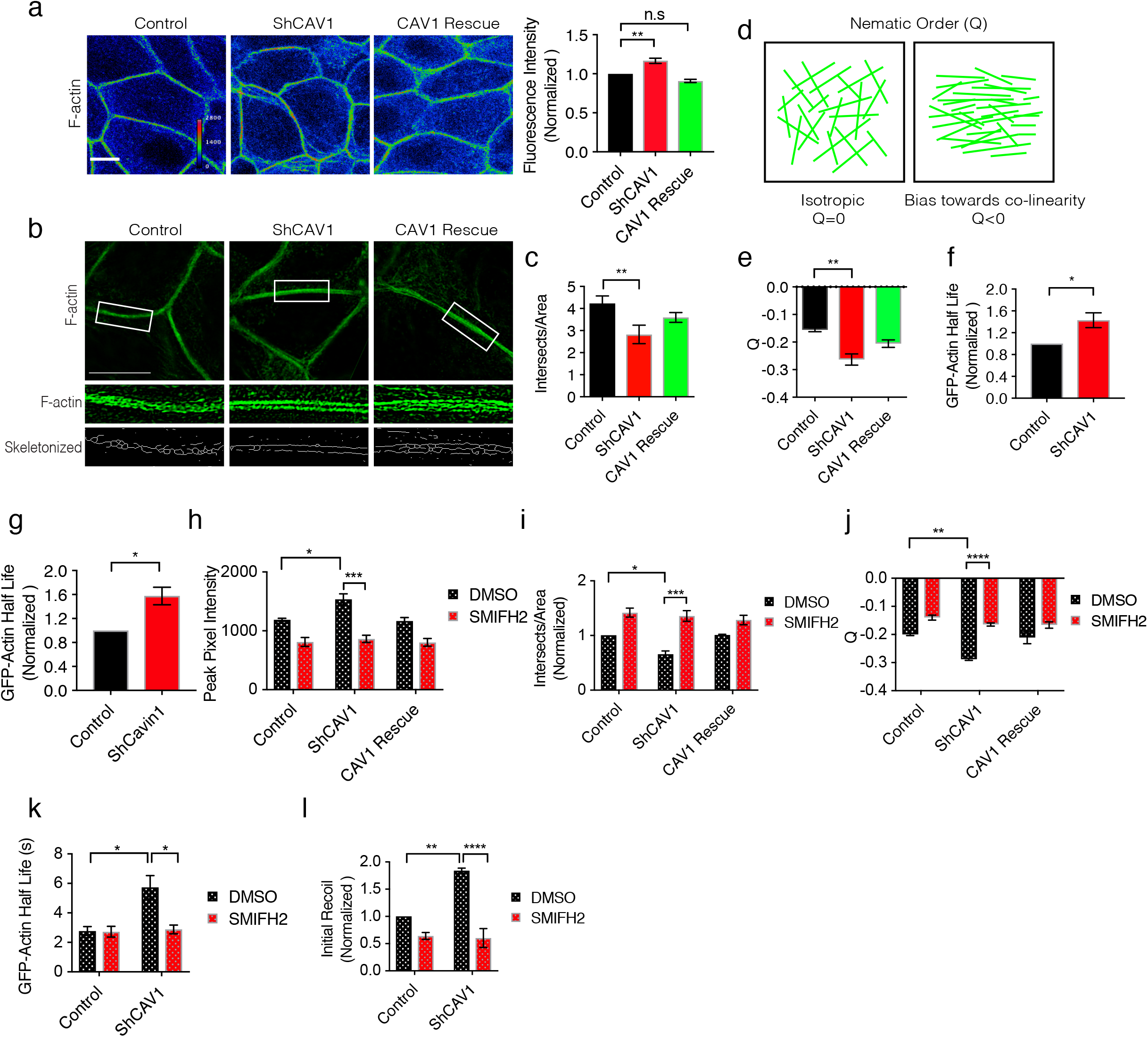
Formin-dependent actin dynamics mediate increased tension in ShCAV1 cells. (a) Effect of CAV1 shRNA on junctional F-actin: representative phalloidin images and quantification of junctional fluorescence intensity. (b,c) Junctional F-actin characterized by SIM and skeletonization analysis. (b) Representative images and quantitation (c). (d,e) Characterization of junctional F-actin organization by nematic order parameter. Schematic diagram of the alignment of actin bundles and nematic order (d) and quantification (e). (f,g) Effect of CAV1 shRNA (f) or Cavin 1 shRNA (g) on junctional F-actin stability: half-life of fluorescence decay after photoactivation of mRFP-PA-GFP-actin at cell junctions. (h-k) Effect of SMIFH2 on the junctional F-actin cytoskeleton in CAV1 KD cells. Quantification of junctional F-actin from phalloidin fluorescence intensity (h); F-actin organization by skeletonization (i) and nematic order analysis (j); and F-actin stability: half-life of fluorescence decay after photoactivation of mRFP-PA-GFP-actin (k). Cells were treated with DMSO (control, 0.1% v/v) or SMIFH2 (20 µM, 30 min). (l) Effect of SMIFH2 on junctional tension in CAV1 KD cells, measured by initial recoil velocity. All data are means ± s.e.m; n.s, not significant; \**P*<0.05, \*\**P*<0.01, \*\*\**P*<0.001, \*\*\*\**P*<0.0001; calculated from N=3 independent experiments analysed with one-way ANOVA Dunnett’s multiple comparisons test except (f) and (g) which were analysed by Student’s *t*-test, (h), (i), (j), (k), (l) which were analysed by two-way ANOVA Tukey’s multiple comparisons test. Scale bar, 10µm.

To pursue this, we first characterized F-actin network architecture by structured illumination microscopy (SIM). Apical junctional F-actin appeared more condensed in CAV1 KD cells (Fig 4b) and fewer overlapping filaments and bundles were evident after skeletonization (Fig 4c). This implied that the co-linear architecture of the junctional cytoskeleton was enhanced by CAV1 depletion. This notion was reinforced by measuring nematic order of junctional F-actin after Fourier transformation of fluorescence signals from the SIM images (Fig 4d) (Reymann et al., 2016). The nematic order tensor (Q) was reduced in CAV1 KD cells (Fig 4e), consistent with decreased isotropy and a more co-linear organization of the actin filaments. Finally, we characterized actin dynamics by expressing G-actin tagged with photoactivatable-GFP (PAGFP-G-actin) and measuring its fluorescence decay after photoactivation (FDAP) at junctions (Fig 4f). The half-life (T_1/12_) of fluorescence decay was significantly increased at the junctions of CAV1 KD (Fig 4f) and Cavin1KD (Fig 4g) cells, compared with controls, implying that the F-actin pool was more stable. Overall, these finding suggested that a mechanism that stabilized F-actin and promoted its bundling at AJ was overactive in CAV1 KD cells.

Formins were attractive candidates to mediate these effects, as they promote the assembly and stabilization of unbranched F-actin networks and are active at cell-cell junctions (Goode and Eck, 2007; Grikscheit and Grosse, 2016; Schonichen and Geyer, 2010). Indeed, the broad-spectrum formin inhibitor SMIFH2 corrected the increased F-actin levels seen at the junctions of CAV1 KD cells (Fig 4h). Further, it also restored F-actin organization (Fig 4i, j) and actin dynamics (Fig 4k) in CAV1 KD cells to levels seen in wild-type or CAV1-reconstituted cells. Finally, SMIFH2 corrected the elevated junctional tension seen in CAV1 KD cells (Fig 4l). This therefore implied that an over-active formin was responsible for increasing junctional tension when CAV1 was depleted.

### FMNL2 formin is upregulated at adherens junctions in CAV1 KD cells

In order to identify the responsible formin(s), we first screened for candidates expressed in AML12 cells with quantitative PCR (Fig S4a). Formin transcript levels were not altered by CAV1 KD (Fig S4b), suggesting that any functional contributions to the CAV1 KD phenotype might arise at a post-transcriptional step. We focused on FMNL2 because its mRNA was relatively abundant (Fig S4a) and FMNL2-EGFP co-localized with E-cadherin at AJ in wild-type cells (a transgene being used in the absence of an antibody that was effective for immunofluorescence, Fig 5a). Furthermore, junctional FMNL2-EGFP was increased by CAV1 KD (Fig 5b; Fig 6b), although this did not alter total expression of endogenous FMNL2 (Fig S4c). Thus, CAV1 deficiency enhanced the junctional recruitment of FMNL2. Consistent with this, the stability of FMNL2-EGFP measured by FRAP was increased at junctions in CAV1 KD cells (Fig 5c,d).

**Figure 5.**
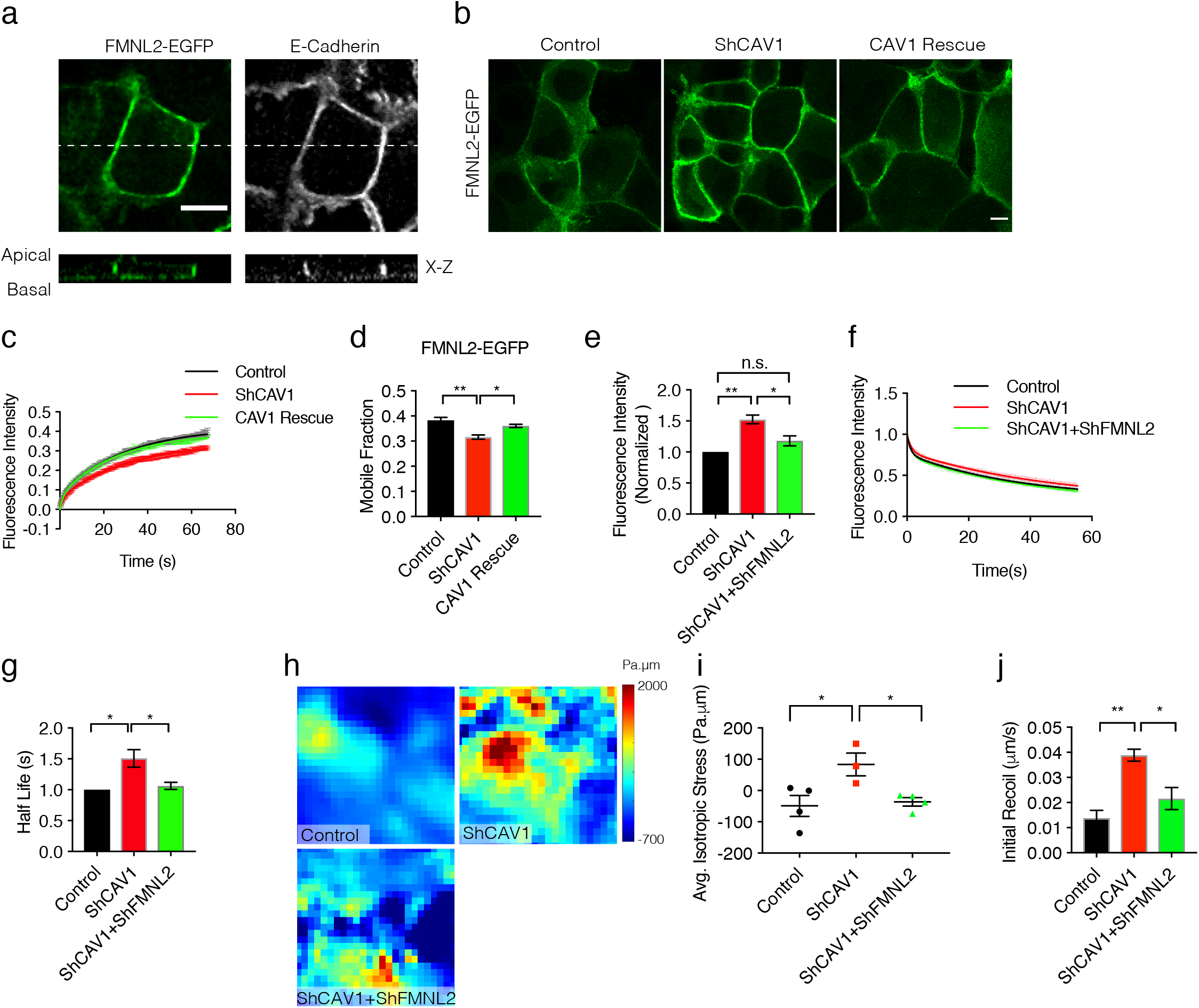
FMNL2 formin is upregulated at adherens junctions in ShCAV1 cells. (a) Representative images of FMNL2-EGFP and endogenous E-cadherin in AML12 cells; respective X-Z views taken at the dotted lines. (b) Representative images of FMNL2-EGFP in GFP control, ShCAV1 GFP and CAV1 Rescue GFP cells. (c,d) Effect of CAV1 KD on the junctional stability of FMNL2-EGFP measured by Fluorescence recovery after photobleaching. (c) Recovery curves and (d) mobile fractions. (e-g) Effect of FMNL2 depletion on the junctional F-actin cytoskeleton in CAV1 shRNA cells. (e) Junctional F-actin from fluorescence intensity of phalloidin staining; and junctional actin stability from fluorescence decay of mRFP-PA-GFP actin: decay curves (f) and half-life (g). (h, i) Effect of FMNL2 depletion on monolayer stress in CAV1 shRNA cells: representative heat maps (h) and (i) quantification of average isotropic stress across 12hrs. (j) Effect of FMNL2 depletion on junctional tension (initial recoil velocity) in CAV1 shRNA cells. All data are means ± s.e.m; n.s, not significant; \**P*<0.05, \*\**P*<0.01, \*\*\**P*<0.001, \*\*\*\**P*<0.0001; calculated from N=3 independent experiments analysed with one-way ANOVA, Tukey’s multiple comparisons test. Scale bar, 10µm.

**Figure 6.**
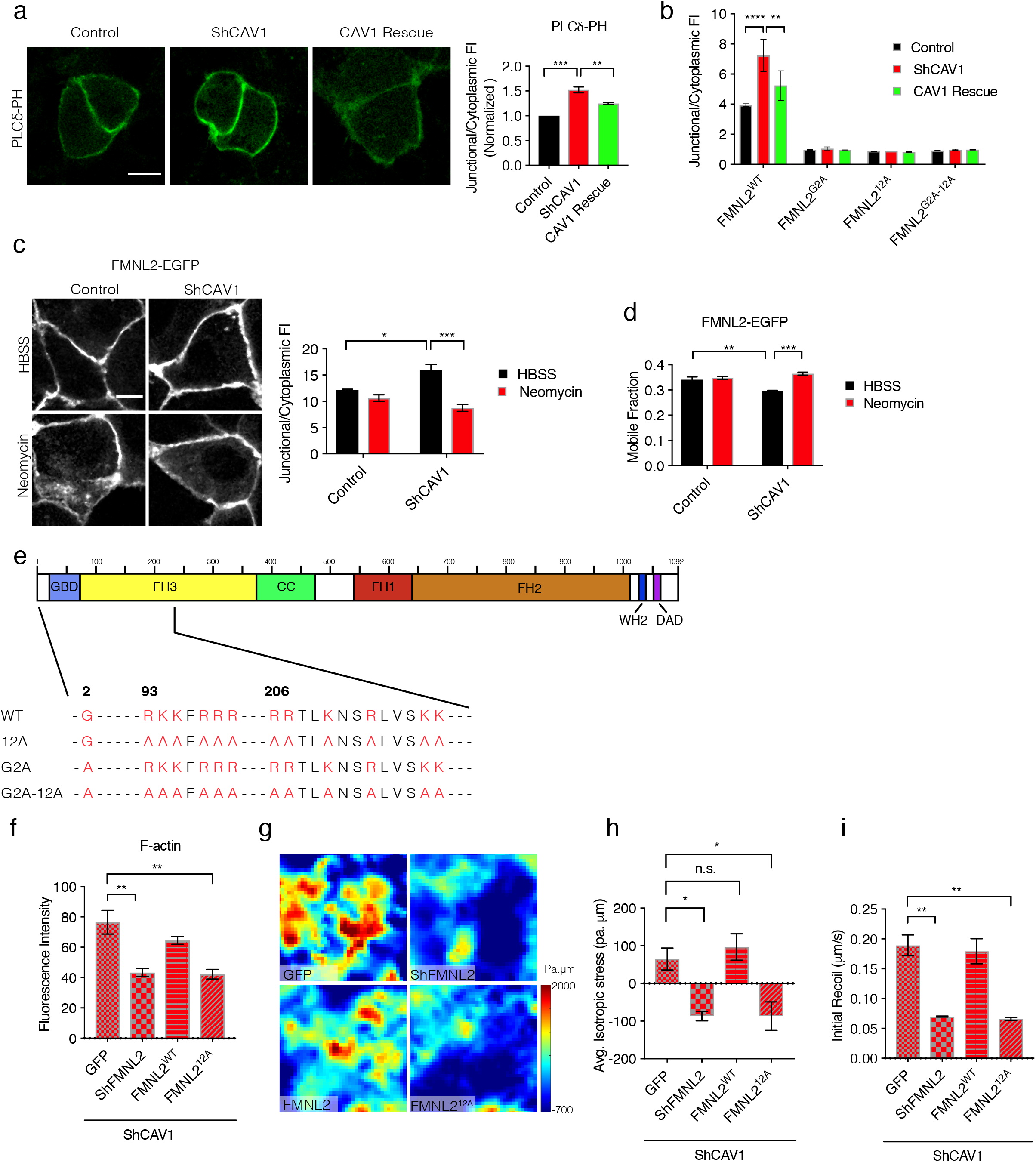
PtdIns(4,5)P_2_ recruits FMNL2 to junctions in ShCAV1 cells. (a) Effect of CAV1 depletion on junctional levels of the PtdIns(4,5)P_2_ location sensor (PLCδ-PH): representative images and quantification. (b) Quantification of junctional FMNL2^WT^, FMNL2^G2A^, FMNL2^12A^ and FMNL2^G2A-12A^ in control, ShCAV1 and CAV1 Rescue cells. (c) Effect of neomycin (3 mM, 1 h) on junctional FMNL2-EGFP levels in control and ShCAV1 cells: representative images and quantitation. (d) Mobile fraction of FMNL2-EGFP calculated from FRAP analysis of control and ShCAV1 with and without neomycin. (e) Schematic diagram of FMNL2 depicting the mutation sites in FMNL2^G2A^, FMNL2^12A^ and FMNL2^G2A-12A^. (f-i) Effect of mutating lipid-binding of FMNL2 on the junctional cytoskeleton and tensile profile of shCAV1 cells. (f) Junctional F-actin levels from fluorescence intensity of phalloidin; monolayer tensile stress from BISM: (g) representative heat maps and (h) average isotropic stress; and (i) initial junctional recoil velocity. shCAV1 cells were further depleted of FMNL2 by shRNA and reconstituted with FMNL2^WT^ or FMNL2^12A^. All data are means ± s.e.m; n.s, not significant; \**P*<0.05, \*\**P*<0.01, \*\*\**P*<0.001, \*\*\*\**P*<0.0001; calculated from N=3 independent experiments analysed with one-way ANOVA, Tukey’s multiple comparisons test except (b), (c) and (d) which were analysed with two-way ANOVA, Tukey’s multiple comparisons test. Scale bar, 10µm.

To test if FMNL2 was responsible for the biomechanical changes that arose from CAV1 deficiency, we depleted FMNL2 in CAV1 KD cells (Fig S4d). FMNL2 shRNA specifically restored F-actin intensity (Fig 5e, Fig 6f) and junctional actin dynamics (Fig 5f,g) of CAV1 KD cells to wild-type levels. Therefore, FMNL2 appeared necessary for the actin cytoskeleton to be dysregulated in CAV1 KD cells. Importantly, FMNL2 KD also specifically corrected the elevated levels of monolayer and junctional tension seen in CAV1 KD monolayers, as measured by BISM (Fig 5h,i) and junctional recoil (Fig 5j, S4e), respectively. Thus, FMNL2 mediated the cytoskeletal changes that increased monolayer tension in CAV1 deficiency. We therefore hypothesized that this might reflect overactivity in the mechanism that recruits FMNL2 to junctions.

### PtdIns(4,5)P_2_ recruits FMNL2 to the junctions of CAV1 KD cells

Multiple signals influence the cortical recruitment of FMNL2, including the Cdc42 GTPase and a number of lipid-binding mechanisms (Block et al., 2012; Kuhn et al., 2015). In particular, FMNL2 contains an N-terminal myristoylation site and a polybasic region in its FH3 domain (Kuhn et al., 2015) that can bind to acidic phospholipids, such as phosphatidylinositol-4,5-bisphosphate (PtdIns(4,5)P_2_). We were unable to detect active Cdc42 at junctions using a location biosensor bearing the WASP GTPase-binding domain (Fig S5a). Therefore, we focused on potential changes in membrane lipids, especially PtdIns(4,5)P_2_, to which cavin can bind (Kovtun et al., 2015; Tillu et al., 2015).

First, we characterized the cellular distribution of PtdIns(4,5)P_2_ using a location biosensor that incorporates the PH domain of phospholipase Cδ (PLCδ-PH). PtdIns(4,5)P_2_ was evident at control AJ and, after correction for expression levels of the sensor, we observed a clear, CAV1-specific, increase in its junctional signal in CAV1 KD (Fig 6a). To test if this might be responsible for recruiting FMNL2, we next blocked access to PtdIns(4,5)P_2_ using neomycin, administered at levels that did not cause evident cell toxicity but displaced PLCδ-PH from junctions (Fig S5b). Adding neomycin to CAV1 KD cells restored both junctional FMNL2 (Fig 6c) and the junctional stability of FMNL2-EGFP (Fig 6d) to control levels. This suggested that increased PtdIns(4,5)P_2_ was responsible for recruiting and stabilizing FMNL2 at the junctions of CAV1 KD cells. We confirmed this with another PtdIns(4,5)P_2_-masking strategy, using PBP10, a cell-permeant peptide derived from the PtdIns(4,5)P_2_-binding site of gelsolin (Fig S5c; (Cunningham et al., 2001)). PBP10 restored junctional FMNL2 in CAV1 KD cells to the same level as control cells (Fig S5e,f), but no effect was seen with a control peptide (RhoB-QRL, Fig S5d, g).

We then refined this analysis by disrupting the lipid-binding motifs of FMNL2 (Fig 6e) (Kuhn et al., 2015). First, we screened the cellular localization of mutant transgenes expressed in FMNL2 shRNA cells. Junctional localization was abolished by mutating the PtdIns(4,5)P_2_-binding capacity of the polybasic domain (FMNL2^12A^; Fig 6e) (Kuhn et al., 2015). The localization of FMNL2^12A^ was completely cytosolic in cells that were wild-type for CAV1 and this was not affected by CAV1 KD (Fig 6b, S5h). This supported the notion that PtdIns(4,5)P_2_ is necessary for cortical recruitment of FMNL2. Interestingly, junctional localization was also consistently abolished when the myristoylation motif was disrupted (FMNL2^G2A^, (Kuhn et al., 2015); Fig6b, S5h) and when both motifs were mutated (FMNL2^G2A-12A^; Fig6b, S5h). Therefore, both the myristoylation and PtdIns(4,5)P_2_-binding mechanisms are likely to participate in its junctional recruitment.

Then we used these mutants to test if membrane recruitment of FMNL2 was specifically required to alter the junctional cytoskeleton in CAV1 KD cells. Consistent with our earlier observations, the elevated junctional F-actin level seen in CAV1 KD cells was corrected by simultaneous FMNL2 KD (Fig 6f). It was restored to an elevated level by reconstitution with FMNL2^WT^, but not by the PtdIns(4,5)P_2_-uncoupled FMNL2^12A^ mutant. Therefore, actin dysregulation in CAV1 KD required junctional FMNL2. Strikingly, junctional FMNL2 was also required for the hyper-contractile phenotype of CAV1 KD. CAV1 KD cells did not show either increased monolayer tension (Fig 6g,h) or increased junctional recoil (Fig 6i, S5j) when endogenous FMNL2 was depleted and reconstituted with FMNL2^12A^. This implies that enhanced junctional recruitment of FMNL2 was responsible for increasing tension in CAV1 KD monolayers.

### Tensional dysregulation by PtdIns(4,5)P_2-_FMNL2 disrupts oncogenic extrusion

Overall, these experiments suggest a pathway to explain how CAV1 KD increases monolayer tension, where increased PtdIns(4,5)P_2_ recruits FMNL2 to stabilize and reorganize F-actin at AJ. Therefore, we interrupted this pathway to test if it was responsible for CAV1 deficient-monolayers being unable to eliminate oncogene-expressing cells.

First, we asked if FMNL2 was necessary for oncogenic extrusion to be compromised in CAV1 KD monolayers. We used cell-mixing experiments where the surrounding CAV1 KD epithelium was further depleted of FMNL2 by shRNA (FMNL2 KD/CAV1 KD cells). We mixed Tet^ON^-HRas^V12^ cells with either FMNL2 KD/CAV1 KD cells (1:20) or CAV1 KD cells for comparison, and grew them to confluence before inducing oncogene expression for 24 h. Strikingly, FMNL2 depletion restored the ability of CAV1 KD cells to extrude HRas^V12^ cells (Fig 7a). Similarly, SMIFH2 restored the ability of CAV1 KD monolayers to extrude HRas^V12^ cells (Fig S6a), supporting the idea that excessive formin activity compromised extrusion in CAV1-depleted monolayers. However, the ability of CAV1-deficient monolayers to extrude HRas^V12^ cells was not restored if the neighbour population was depleted of either FMNL3 or INF2 (Fig 7a, S4d). This reinforced the notion that FMNL2 was selectively responsible for disrupting extrusion in CAV1-deficient monolayers.

**Figure 7.**
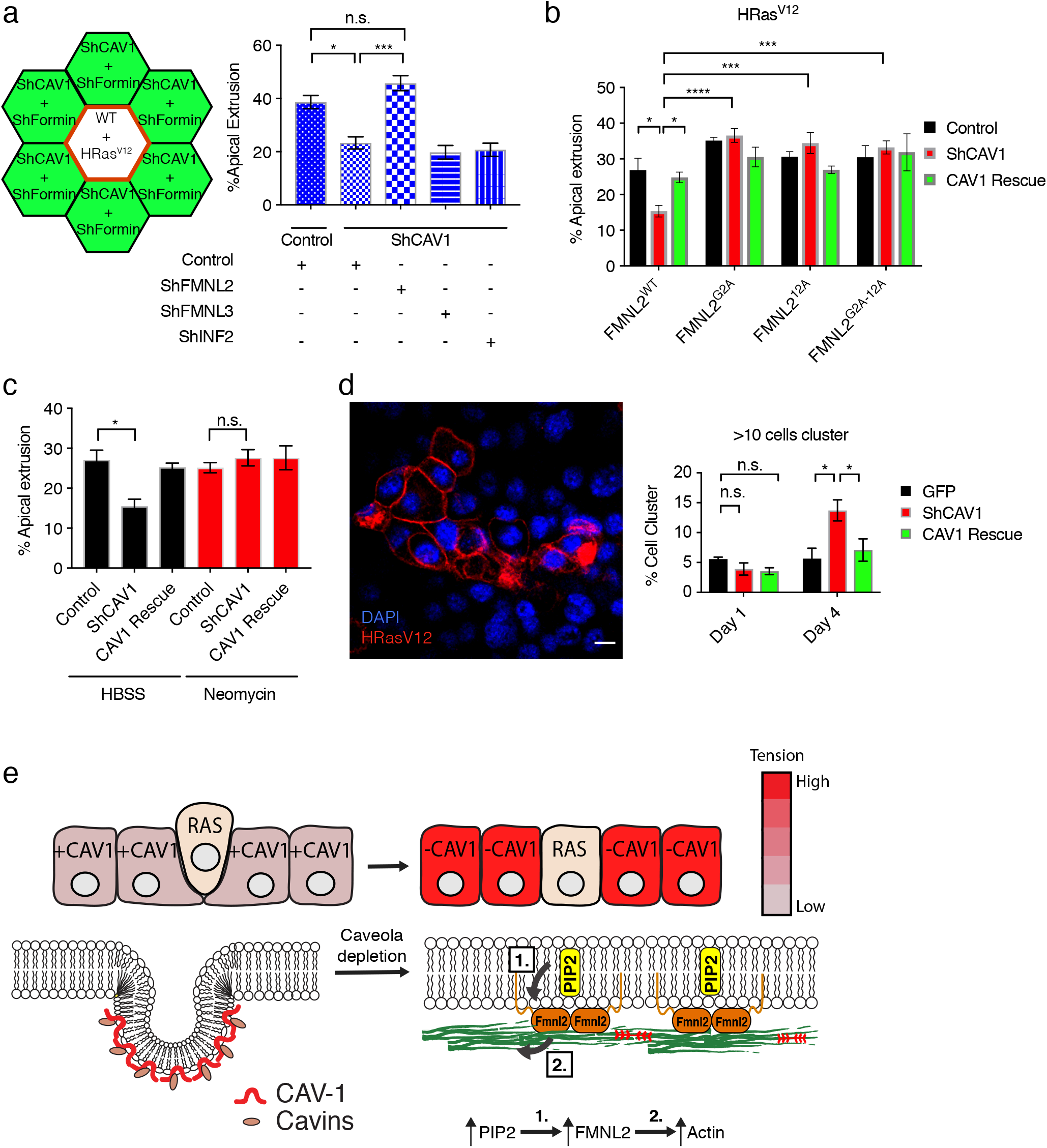
The PtdIns(4,5)P_2_-FMNL2 pathway disrupts oncogenic extrusion in ShCAV1 monolayers. (a) Effect on apical oncogenic extrusion when FMNL2 was depleted in shCAV1 epithelium. Tet^on^ HRas^V12^-expressing WT cells were mixed with a majority of ShCAV1 cells downregulated for FMNL2, FMNL3 or INF2. Schematic illustration of the experimental design and quantification of apical cell extrusion. (b) Disabling lipid-binding of FMNL2 restores oncogenic extrusion to shCAV1 monolayers. Tet^on^ HRas^V12^-expressing WT cells were surrounded by control, ShCAV1 and CAV1 Rescue cells that were further depleted of endogenous FMNL2 and reconstituted with FMNL2^WT^ FMNL2^G2A^, FMNL2^12A^ or FMNL2^G2A-12A^. (c) Apical oncogenic cell extrusion of Tet^on^ HRas^V12^-expressing WT cells mixed with GFP control, ShCAV1 GFP and RescueCAV1 GFP cells with and without neomycin treatment. (d) Oncogene-expressing cells form clusters when apical extrusion is disrupted by CAV1 depletion. Tet^on^ HRas^V12^ expressed in WT cells were mixed with ShCAV1 neighbours. Representative image after 4 days. Clusters containing > 10 transgene-expressing cells are expressed as a percentage of all clusters (see Methods for details). (e) Schematic diagram: Apical extrusion of oncogene-expressing cells is inhibited by elevated monolayer tension upon CAV1 depletion. Caveola depletion elevates monolayer tension by increasing Ptdlns(4,5)P_2_ at the plasma membrane. This recruits FMNL2 (1) to stabilize cortical F-actin (2). All data are means ± s.e.m; n.s, not significant; \**P*<0.05, \*\**P*<0.01, \*\*\**P*<0.001, \*\*\*\**P*<0.0001; calculated from N=3 independent experiments. (a) and (c) were analysed with one-way ANOVA, Tukey’s multiple comparisons test, (b) and (d) were analysed with two-way ANOVA, Tukey’s multiple comparisons test. Scale bar, 10µm.

We then asked if junctional recruitment of FMNL2 by PtdIns(4,5)P_2_ was responsible for compromising extrusion in CAV1 KD cells. For this, we reconstituted the FMNL2 KD/CAV1 KD neighbour cell population with lipid-binding FMNL2 mutants (Fig 7b). Oncogenic extrusion was compromised when we reconstituted the surrounding epithelium with wild-type FMNL2 but not with FMNL2^12A^ (Fig 7b), suggesting that PtdIns(4,5)P_2_-binding was necessary for FMNL2 to inhibit extrusion in CAV1 KD cells. Consistent with this, the capacity to extrude HRas^V12^ cells was also restored to CAV1 KD cells when the accessibility of PtdIns(4,5)P_2_ was blocked with neomycin (Fig 7c). Together, these argue that recruitment of FMNL2 by increased PtdIns(4,5)P_2_ participates in disrupting extrusion in CAV1-deficient cells. Oncogenic extrusion was also restored when FMNL2 KD/CAV1 KD neighbour cells were reconstituted with either FMNL2^G2A^ or FMNL2^G2A-12A^ (Fig7b). These findings reinforce the concept that cortical recruitment of FMNL2 was necessary to disrupt extrusion in CAV1-deficient cells.

### Retained oncogene cells proliferate when extrusion is disrupted by CAV1 KD

Finally, we asked what was the fate of oncogene-expressing cells when extrusion was disrupted (Fig 7d, S6b). After mixing Tet^ON^-HRas^V12^ cells with either WT or CAV1 KD AML12 cells, we induced H-Ras^V12^ for 4 days and counted the number of large (>10 cell) clusters of H-Ras^V12^-positive cells present within the monolayer. Relatively few such clusters were seen when the neighbouring epithelium was wild-type. However, the incidence of these clusters was significantly increased when extrusion was blocked with CAV1 KD in the neighbour cells (Fig 7d). The effect of CAV1 KD was reversed by expression of exogenous CAV1, confirming its specificity (Fig 7d). These results strongly suggest that oncogene-expressing cells were retained in epithelia and proliferated when CAV1-deficient neighbours were unable to eliminate them by apical extrusion.

## Discussion

Caveolae are mechanosensitive organelles that can influence cellular mechanics. Our findings now reveal how caveolae also affect tissue mechanics to support epithelial homeostasis. They lead to two conclusions. First, caveolae can set tissue-scale levels of contractile tension by a lipid-based signalling pathway that regulates the cortical F-actin cytoskeleton. Second, this affects the ability of epithelia to eliminate oncogene-expressing cells by apical extrusion. It carries the implication that tissue mechanics can complement cell-autonomous changes in the tumor cell to influence cancer outcome.

The intrinsic tissue tension of epithelial monolayers arises from, and is propagated by, the AJ that couple the contractile cortices of cells together (Charras and Yap, 2018). CAV1 depletion increased epithelial monolayer tension in our experiments, evident at length scales from molecular-level tension to tissue-scale stress. Although CAV1 can have caveola-independent effects (Hayer et al., 2010b; Hill et al., 2008), cavin-1 depletion also reduced junctional tension without affecting CAV1 levels. This common effect suggested that changes in caveola abundance can set levels of tissue tension in monolayers. Changes in cellular contractility are commonly thought to arise when signals such as MLCK and RhoA-ROCK alter the activity of Myosin II (Heer and Martin, 2017). However, neither the level nor activity of Myosin II was detectably changed in CAV1 KD cells. Instead, the enhanced contractile tension was attributable to increased levels of PtdIns(4,5)P_2_ which recruited FMNL2 to the junctional membrane. Thus, the hypercontractile phenotype of CAV1-depleted cells could be corrected by blocking protein access to PtdIns(4,5)P_2_ or depleting FMNL2 itself. With this, junctional F-actin dynamics and organization were restored to wild-type levels, consistent with the actin-regulatory effects of FMNL2 (Block et al., 2012; Grikscheit et al., 2015; Grikscheit and Grosse, 2016).

FMNL2 (and its related FMNL3) are unusual amongst the formins because they bear multiple lipid-binding domains (Kuhn et al., 2015). These domains proved to be critical in our experiments, as contractility could be restored to normal by mutating the ability of FMNL2 to bind PtdIns(4,5)P_2_. Together, this indicated that FMNL2 was being recruited in response to the elevated levels of PtdIns(4,5)P_2_. This mechanism appeared to cooperate with the N-terminal myristoylation site, which is necessary, but not sufficient, for FMNL2 to interact stably with the membrane (Block et al., 2012). Interestingly, although FMNL2 can also respond to Cdc42, we were unable to detect junctional Cdc42 activity in our experiments, consistent with earlier evidence that lipid-binding alone could support membrane recruitment of FMNL2 (Block et al., 2012). Therefore, the increased levels of PtdIns(4,5)P_2_ appeared to be responsible for increasing monolayer tension when caveolae were depleted.

PtdIns(4,5)P_2_ has been observed to concentrate in caveolae (Bucki et al., 2019; Fujita et al., 2009; Parton and Collins, 2016) and cavins can bind PtdIns(4,5)P_2_ directly (Tillu et al., 2015). Thus, caveolar downregulation may have released this pool of PtdIns(4,5)P_2_ into the bulk membrane for signaling. Caveolae can also influence membrane organization at the nanoscale (Ariotti et al., 2014), with the potential of localizing PtdIns(4,5)P_2_ in regions of the membrane. Irrespective of the precise mechanism, these findings reveal an alternative pathway for lipid signalling to control cell contractility by regulating actin dynamics and organization. Since tension was also reduced when Myosin II was inhibited with Y-27632 or blebbistatin (not shown), the PtdIns(4,5)P_2_-FMNL2 pathway is likely to complement the canonical activators of Myosin II by modulating the actin scaffolds that are also necessary for force generation (Linsmeier et al., 2016; Murrell and Gardel, 2012).

Upregulation of the PtdIns(4,5)P_2_-FMNL2 pathway coincided with a defect in the ability of CAV1 KD cells to eliminate oncogene-transfected cells by apical extrusion. Oncogenic extrusion entails an active response of the host epithelium, presumably responding to signals from the oncogene-expressing cell (Hogan et al., 2009; Porazinski et al., 2016). Therefore, it was striking to find that caveolae were required only in the surrounding epithelium for the extrusion response to occur. Extrusion occurred normally when CAV1 was depleted in the oncogene-expressing cell, but was defective when CAV1 was depleted in the epithelium. This suggested that the defect in oncogenic extrusion reflected some general change in CAV1-depleted monolayers, rather than a change in the transformed cells themselves. Importantly, extrusion was restored to CAV1 KD epithelia by depleting FMNL2 or by interrupting its recruitment by PtdIns(4,5)P_2_. As these were exactly the maneuvers that restored tissue tension levels to normal, it suggested that extrusion was being disrupted by the general increase in tissue tension that the overactive PtdIns(4,5)P_2_-FMNL2 pathway elicited. Alternatively, it was possible that caveolae participated in allowing neighbour cells to detect the nearby presence of oncogene-expressing cells (Hogan et al., 2009). However, extrusion was also blocked in CAV1 wild-type epithelia when tension was increased by expression of exogenous MRLC. This reinforced the notion that altered tissue tension was key to the defect in oncogenic extrusion.

How might increased tissue tension inhibit the ability of the epithelium to extrude transformed cells? Contractile tension is focused in the zonulae adherente of polarized epithelial cells, and this contributes to retaining cells within monolayers (Wu et al., 2014). Indeed, oncogenic extrusion was associated with downregulation of tension at the apical-lateral interface (Wu et al., 2014). Therefore, re-setting tissue tension to a higher level, either indirectly by disrupting caveolae or directly by expressing MRLC, may have prevented apical tension from being reduced to a level sufficient to permit extrusion. As well, tissue tension can influence processes such as cell-cell rearrangements that may be necessary for epithelia to accommodate the elimination of cells (Fernandez-Gonzalez et al., 2009; Kobb et al., 2017; Marinari et al., 2012; Rauzi et al., 2008; Samarage et al., 2015). Irrespective of the detailed mechanism, our findings argue that re-setting of tissue tension by the PtdIns(4,5)P_2_-FMNL2 pathway disrupted the homeostatic response to transformation in CAV1 KD cells.

Many studies have sought to characterize how caveolae influence cancer; these have revealed complex links between the expression of caveolar components and tumour progression or suppression, which have been difficult to resolve. Typically, these experiments have been interpreted from the cell-autonomous perspective of the tumor cell (Engelman et al., 1997; Lee et al., 1998; Sagara et al., 2004; Yang et al., 1998). Our results suggest that it is also important to consider the caveola-status of the host epithelium. Apical extrusion has been proposed to represent an early-response mechanism that allows epithelia to eliminate newly-transformed cells from the body, potentially before they have proliferated to any significant degree (Hogan et al., 2009; Kajita and Fujita, 2015; Kajita et al., 2010; Kon et al., 2017; Ohoka et al., 2015; Sasaki et al., 2018). Failure of the epithelium to execute oncogenic extrusion would then be predicted to promote tumor cell retention and potential proliferation. Indeed, this is what we observed when CAV1 was depleted in neighbour cells, even though the oncogene-expressing cells were CAV1-wildtype. This may be one pathway for CAV1 deficiency to prime epithelia, increasing their susceptibility to DMBA-induced tumorigenesis (Capozza et al., 2003). Recently, the caveolar status of cancer-associated fibroblasts was reported to influence tumor development (Goetz et al., 2011). Our observations imply that the host epithelium should also be considered as part of the tumor microenvironment.

We conclude that caveolar status influences epithelial mechanics and homeostasis by controlling the levels of PtdIns(4,5)P_2_ that are available to signal at the membrane. While our experiments utilized stable depletion of caveolar components, membrane lipid composition and organization changed in similar fashion when caveolae were dynamically disassembled in response to hypotonic stress as when they were stably disrupted by CAV1 or cavin KD (Ariotti et al., 2014). It will therefore be interesting to test if the PtdIns(4,5)P_2_-FMNL2 pathway is also engaged when caveolae undergo dynamic disassembly. Such disassembly occurs when membrane tension increases in response to mechanical stresses. Rosettes or clusters of caveolae are especially sensitive to caveola flattening (Golani et al., 2019; Lo et al., 2016; Lo et al., 2015; Sinha et al., 2011; Yeow et al., 2017) and were a particular feature of the caveolae in the junctional regions of the epithelial cells studied here. Thus, caveolae may complement other mechanotransduction pathways found at cell-cell junctions by conferring sensitivity to changes in membrane tension. More generally, our findings highlight the capacity for PtdIns(4,5)P_2_ to control epithelial mechanics. Other mechanisms exist at junctions to influence PtdIns(4,5)P_2_, including lipid kinases and phosphatases as well as lipid-binding scaffolds (Balla, 2013; Budnar et al., 2019; Laux et al., 2000). Caveolae may then collaborate with a regulatory network that tunes PtdIns(4,5)P_2_ to set the level of tensional stress in monolayers.

## Acknowledgements

We thank Philippe Marcq (Institut Curie) for his help in implementing the BISM analysis; our colleagues who provided generous gifts of reagents; and our lab members for their support and good counsel. This work was supported by the National Health and Medical Research Council of Australia (grants 1140090 to R.G.P. and A.Y.; APP1140064 and APP1150083 and fellowship APP1156489 to R.G.P.; and fellowship APP1139592 to AY). RGP is supported by the Australian Research Council (ARC) Centre of Excellence in Convergent Bio-Nano Science and Technology. J.L.T. was the recipient of an Equity Trustees PhD Scholarship in Medical Research; IN by the European Molecular Biological Organization (EMBO ALTF 251-2018); HKK by a post-doctoral fellowship from The Uehara Memorial Foundation; SJS by a project grant from the Rebecca L. Cooper Medical Research Foundation (PG2018168) and UQECR award 1946705; and LB and BL by European Research Council (Grant No. CoG-617233), the LABEX “Who Am I?,” and the Agence Nationale de la Recherche “POLCAM” (Grant No. ANR-17-CE13-0013).

## Author contributions

JLT, RGP and ASY conceived the project; JLT principally conducted the experiments and analysed the data with support from SV, VT, RT (FIB EM), RJJ, and IN (deconvolution confocal imaging); BA developed the α-CAT-TS sensor; GAG, KAM, HKK, and SJS provided guidance and input; LB and BL supported the BISM; and JLT, RGP and ASY wrote the MS.

## References

Acharya, B.R., Nestor-Bergmann, A., Liang, X., Gupta, S., Duszyc, K., Gauquelin, E., Gomez, G.A., Budnar, S., Marcq, P., Jensen, O.E., et al. (2018). A Mechanosensitive RhoA Pathway that Protects Epithelia against Acute Tensile Stress. Dev Cell 47, 439–452 e436.

Acharya, B.R., Wu, S.K., Lieu, Z.Z., Parton, R.G., Grill, S.W., Bershadsky, A.D., Gomez, G.A., and Yap, A.S. (2017). Mammalian Diaphanous 1 Mediates a Pathway for E-cadherin to Stabilize Epithelial Barriers through Junctional Contractility. Cell Rep 18, 2854–2867.

Ariotti, N., Fernandez-Rojo, M.A., Zhou, Y., Hill, M.M., Rodkey, T.L., Inder, K.L., Tanner, L.B., Wenk, M.R., Hancock, J.F., and Parton, R.G. (2014). Caveolae regulate the nanoscale organization of the plasma membrane to remotely control Ras signaling. J Cell Biol 204, 777–792.

Balla, T. (2013). Phosphoinositides: tiny lipids with giant impact on cell regulation. Physiol Rev 93, 1019–1137.

Block, J., Breitsprecher, D., Kuhn, S., Winterhoff, M., Kage, F., Geffers, R., Duwe, P., Rohn, J.L., Baum, B., Brakebusch, C., et al. (2012). FMNL2 drives actin-based protrusion and migration downstream of Cdc42. Curr Biol 22, 1005–1012.

Bucki, R., Wang, Y.H., Yang, C., Kandy, S.K., Fatunmbi, O., Bradley, R., Pogoda, K., Svitkina, T., Radhakrishnan, R., and Janmey, P.A. (2019). Lateral distribution of phosphatidylinositol 4,5-bisphosphate in membranes regulates formin- and ARP2/3-mediated actin nucleation. J Biol Chem 294, 4704–4722.

Budnar, S., Husain, K.B., Gomez, G.A., Naghibosidat, M., Varma, A., Verma, S., Hamilton, N.A., Morris, R.G., and Yap, A.S. (2019). Anillin promotes cell contractility by cyclic resetting of RhoA residence kinetics.. Dev Cell In Press.

Capozza, F., Williams, T.M., Schubert, W., McClain, S., Bouzahzah, B., Sotgia, F., and Lisanti, M.P. (2003). Absence of caveolin-1 sensitizes mouse skin to carcinogen-induced epidermal hyperplasia and tumor formation. Am J Pathol 162, 2029–2039.

Charras, G., and Yap, A.S. (2018). Tensile Forces and Mechanotransduction at Cell-Cell Junctions. Curr Biol 28, R445–R457.

Chugh, P., and Paluch, E.K. (2018). The actin cortex at a glance. J Cell Sci 131.

Cunningham, C.C., Vegners, R., Bucki, R., Funaki, M., Korde, N., Hartwig, J.H., Stossel, T.P., and Janmey, P.A. (2001). Cell permeant polyphosphoinositide-binding peptides that block cell motility and actin assembly. J Biol Chem 276, 43390–43399.

Engelman, J.A., Wykoff, C.C., Yasuhara, S., Song, K.S., Okamoto, T., and Lisanti, M.P. (1997). Recombinant expression of caveolin-1 in oncogenically transformed cells abrogates anchorage-independent growth. J Biol Chem 272, 16374–16381.

Fanning, A.S., Van Itallie, C.M., and Anderson, J.M. (2012). Zonula occludens-1 and −2 regulate apical cell structure and the zonula adherens cytoskeleton in polarized epithelia. Mol Biol Cell 23, 577–590.

Fernandez-Gonzalez, R., Simoes Sde, M., Roper, J.C., Eaton, S., and Zallen, J.A. (2009). Myosin II dynamics are regulated by tension in intercalating cells. Dev Cell 17, 736–743.

Fujita, A., Cheng, J., Tauchi-Sato, K., Takenawa, T., and Fujimoto, T. (2009). A distinct pool of phosphatidylinositol 4,5-bisphosphate in caveolae revealed by a nanoscale labeling technique. Proc Natl Acad Sci U S A 106, 9256–9261.

Goetz, J.G., Minguet, S., Navarro-Lerida, I., Lazcano, J.J., Samaniego, R., Calvo, E., Tello, M., Osteso-Ibanez, T., Pellinen, T., Echarri, A., et al. (2011). Biomechanical remodeling of the microenvironment by stromal caveolin-1 favors tumor invasion and metastasis. Cell 146, 148–163.

Golani, G., Ariotti, N., Parton, R.G., and Kozlov, M.M. (2019). Membrane Curvature and Tension Control the Formation and Collapse of Caveolar Superstructures. Dev Cell 48, 523–538 e524.

Goode, B.L., and Eck, M.J. (2007). Mechanism and function of formins in the control of actin assembly. Annu Rev Biochem 76, 593–627.

Grikscheit, K., Frank, T., Wang, Y., and Grosse, R. (2015). Junctional actin assembly is mediated by Formin-like 2 downstream of Rac1. J Cell Biol 209, 367–376.

Grikscheit, K., and Grosse, R. (2016). Formins at the Junction. Trends Biochem Sci 41, 148–159.

Hayer, A., Stoeber, M., Bissig, C., and Helenius, A. (2010a). Biogenesis of caveolae: stepwise assembly of large caveolin and cavin complexes. Traffic 11, 361–382.

Hayer, A., Stoeber, M., Ritz, D., Engel, S., Meyer, H.H., and Helenius, A. (2010b). Caveolin-1 is ubiquitinated and targeted to intralumenal vesicles in endolysosomes for degradation. J Cell Biol 191, 615–629.

Heer, N.C., and Martin, A.C. (2017). Tension, contraction and tissue morphogenesis. Development 144, 4249–4260.

Hill, M.M., Bastiani, M., Luetterforst, R., Kirkham, M., Kirkham, A., Nixon, S.J., Walser, P., Abankwa, D., Oorschot, V.M., Martin, S., et al. (2008). PTRF-Cavin, a conserved cytoplasmic protein required for caveola formation and function. Cell 132, 113–124.

Hogan, C., Dupre-Crochet, S., Norman, M., Kajita, M., Zimmermann, C., Pelling, A.E., Piddini, E., Baena-Lopez, L.A., Vincent, J.P., Itoh, Y., et al. (2009). Characterization of the interface between normal and transformed epithelial cells. Nat Cell Biol 11, 460–467.

Kajita, M., and Fujita, Y. (2015). EDAC: Epithelial defence against cancer-cell competition between normal and transformed epithelial cells in mammals. J Biochem 158, 15–23.

Kajita, M., Hogan, C., Harris, A.R., Dupre-Crochet, S., Itasaki, N., Kawakami, K., Charras, G., Tada, M., and Fujita, Y. (2010). Interaction with surrounding normal epithelial cells influences signalling pathways and behaviour of Src-transformed cells. J Cell Sci 123, 171–180.

Kajita, M., Sugimura, K., Ohoka, A., Burden, J., Suganuma, H., Ikegawa, M., Shimada, T., Kitamura, T., Shindoh, M., Ishikawa, S., et al. (2014). Filamin acts as a key regulator in epithelial defence against transformed cells. Nat Commun 5, 4428.

Kobb, A.B., Zulueta-Coarasa, T., and Fernandez-Gonzalez, R. (2017). Tension regulates myosin dynamics during Drosophila embryonic wound repair. J Cell Sci 130, 689–696.

Kon, S., Ishibashi, K., Katoh, H., Kitamoto, S., Shirai, T., Tanaka, S., Kajita, M., Ishikawa, S., Yamauchi, H., Yako, Y., et al. (2017). Cell competition with normal epithelial cells promotes apical extrusion of transformed cells through metabolic changes. Nat Cell Biol 19, 530–541.

Kovacs, E.M., Goodwin, M., Ali, R.G., Paterson, A.D., and Yap, A.S. (2002). Cadherin-directed actin assembly: E-cadherin physically associates with the Arp2/3 complex to direct actin assembly in nascent adhesive contacts. Curr Biol 12, 379–382.

Kovtun, O., Tillu, V.A., Ariotti, N., Parton, R.G., and Collins, B.M. (2015). Cavin family proteins and the assembly of caveolae. J Cell Sci 128, 1269–1278.

Kuhn, S., Erdmann, C., Kage, F., Block, J., Schwenkmezger, L., Steffen, A., Rottner, K., and Geyer, M. (2015). The structure of FMNL2-Cdc42 yields insights into the mechanism of lamellipodia and filopodia formation. Nat Commun 6, 7088.

Laux, T., Fukami, K., Thelen, M., Golub, T., Frey, D., and Caroni, P. (2000). GAP43, MARCKS, and CAP23 modulate PI(4,5)P_2_ at plasmalemmal rafts, and regulate cell cortex actin dynamics through a common mechanism. J Cell Biol 149, 1455–1471.

le Duc, Q., Shi, Q., Blonk, I., Sonnenberg, A., Wang, N., Leckband, D., and de Rooij, J. (2010). Vinculin potentiates E-cadherin mechanosensing and is recruited to actin-anchored sites within adherens junctions in a myosin II-dependent manner. J Cell Biol 189, 1107–1115.

Lee, S.W., Reimer, C.L., Oh, P., Campbell, D.B., and Schnitzer, J.E. (1998). Tumor cell growth inhibition by caveolin re-expression in human breast cancer cells. Oncogene 16, 1391–1397.

Leung, C.T., and Brugge, J.S. (2012). Outgrowth of single oncogene-expressing cells from suppressive epithelial environments. Nature 482, 410–413.

Linsmeier, I., Banerjee, S., Oakes, P.W., Jung, W., Kim, T., and Murrell, M.P. (2016). Disordered actomyosin networks are sufficient to produce cooperative and telescopic contractility. Nature Communications 7, 12615.

Liu, L., Brown, D., McKee, M., Lebrasseur, N.K., Yang, D., Albrecht, K.H., Ravid, K., and Pilch, P.F. (2008). Deletion of Cavin/PTRF causes global loss of caveolae, dyslipidemia, and glucose intolerance. Cell Metab 8, 310–317.

Lo, H.P., Hall, T.E., and Parton, R.G. (2016). Mechanoprotection by skeletal muscle caveolae. Bioarchitecture 6, 22–27.

Lo, H.P., Nixon, S.J., Hall, T.E., Cowling, B.S., Ferguson, C., Morgan, G.P., Schieber, N.L., Fernandez-Rojo, M.A., Bastiani, M., Floetenmeyer, M., et al. (2015). The caveolin-cavin system plays a conserved and critical role in mechanoprotection of skeletal muscle. J Cell Biol 210, 833–849.

Marinari, E., Mehonic, A., Curran, S., Gale, J., Duke, T., and Baum, B. (2012). Live-cell delamination counterbalances epithelial growth to limit tissue overcrowding. Nature 484, 542–545.

Martin, A.C., Gelbart, M., Fernandez-Gonzalez, R., Kaschube, M., and Wieschaus, E.F. (2010). Integration of contractile forces during tissue invagination. J Cell Biol 188, 735–749.

Murrell, M.P., and Gardel, M.L. (2012). F-actin buckling coordinates contractility and severing in a biomimetic actomyosin cortex. Proc Natl Acad Sci U S A 109, 20820–20825.

Nier, V., Jain, S., Lim, C.T., Ishihara, S., Ladoux, B., and Marcq, P. (2016). Inference of Internal Stress in a Cell Monolayer. Biophys J 110, 1625–1635.

Ohoka, A., Kajita, M., Ikenouchi, J., Yako, Y., Kitamoto, S., Kon, S., Ikegawa, M., Shimada, T., Ishikawa, S., and Fujita, Y. (2015). EPLIN is a crucial regulator for extrusion of RasV12-transformed cells. J Cell Sci 128, 781–789.

Parton, R.G., and Collins, B.M. (2016). Unraveling the architecture of caveolae. Proc Natl Acad Sci U S A 113, 14170–14172.

Parton, R.G., and del Pozo, M.A. (2013). Caveolae as plasma membrane sensors, protectors and organizers. Nat Rev Mol Cell Biol 14, 98–112.

Porazinski, S., de Navascues, J., Yako, Y., Hill, W., Jones, M.R., Maddison, R., Fujita, Y., and Hogan, C. (2016). EphA2 Drives the Segregation of Ras-Transformed Epithelial Cells from Normal Neighbors. Curr Biol 26, 3220–3229.

Ratheesh, A., Gomez, G.A., Priya, R., Verma, S., Kovacs, E.M., Jiang, K., Brown, N.H., Akhmanova, A., Stehbens, S.J., and Yap, A.S. (2012). Centralspindlin and alpha-catenin regulate Rho signalling at the epithelial zonula adherens. Nat Cell Biol 14, 818–828.

Rauzi, M., Krzic, U., Saunders, T.E., Krajnc, M., Ziherl, P., Hufnagel, L., and Leptin, M. (2015). Embryo-scale tissue mechanics during Drosophila gastrulation movements. Nat Commun 6, 8677.

Rauzi, M., Verant, P., Lecuit, T., and Lenne, P.F. (2008). Nature and anisotropy of cortical forces orienting Drosophila tissue morphogenesis. Nat Cell Biol 10, 1401–1410.

Reymann, A.C., Staniscia, F., Erzberger, A., Salbreux, G., and Grill, S.W. (2016). Cortical flow aligns actin filaments to form a furrow. Elife 5.

Sagara, Y., Mimori, K., Yoshinaga, K., Tanaka, F., Nishida, K., Ohno, S., Inoue, H., and Mori, M. (2004). Clinical significance of Caveolin-1, Caveolin-2 and HER2/neu mRNA expression in human breast cancer. Br J Cancer 91, 959–965.

Samarage, C.R., White, M.D., Alvarez, Y.D., Fierro-Gonzalez, J.C., Henon, Y., Jesudason, E.C., Bissiere, S., Fouras, A., and Plachta, N. (2015). Cortical Tension Allocates the First Inner Cells of the Mammalian Embryo. Dev Cell 34, 435–447.

Sasaki, A., Nagatake, T., Egami, R., Gu, G., Takigawa, I., Ikeda, W., Nakatani, T., Kunisawa, J., and Fujita, Y. (2018). Obesity Suppresses Cell-Competition-Mediated Apical Elimination of RasV12-Transformed Cells from Epithelial Tissues. Cell Rep 23, 974–982.

Saw, T.B., Doostmohammadi, A., Nier, V., Kocgozlu, L., Thampi, S., Toyama, Y., Marcq, P., Lim, C.T., Yeomans, J.M., and Ladoux, B. (2017). Topological defects in epithelia govern cell death and extrusion. Nature 544, 212–216.

Schonichen, A., and Geyer, M. (2010). Fifteen formins for an actin filament: a molecular view on the regulation of human formins. Biochim Biophys Acta 1803, 152–163.

Sinha, B., Koster, D., Ruez, R., Gonnord, P., Bastiani, M., Abankwa, D., Stan, R.V., Butler-Browne, G., Vedie, B., Johannes, L., et al. (2011). Cells respond to mechanical stress by rapid disassembly of caveolae. Cell 144, 402–413.

Tillu, V.A., Kovtun, O., McMahon, K.A., Collins, B.M., and Parton, R.G. (2015). A phosphoinositide-binding cluster in cavin1 acts as a molecular sensor for cavin1 degradation. Mol Biol Cell 26, 3561–3569.

Trepat, X., and Sahai, E. (2018). Mesoscale physical principles of collective cell organization. Nature Physics 14, 671–682.

Trepat, X., Wasserman, M.R., Angelini, T.E., Millet, E., Weitz, D.A., Butler, J.P., and Fredberg, J.J. (2009). Physical forces during collective cell migration. Nature Physics 5, 426.

Tsukita, S., Katsuno, T., Yamazaki, Y., Umeda, K., Tamura, A., and Tsukita, S. (2009). Roles of ZO-1 and ZO-2 in establishment of the belt-like adherens and tight junctions with paracellular permselective barrier function. Ann N Y Acad Sci 1165, 44–52.

Turner, J.R., Rill, B.K., Carlson, S.L., Carnes, D., Kerner, R., Mrsny, R.J., and Madara, J.L. (1997). Physiological regulation of epithelial tight junctions is associated with myosin light-chain phosphorylation. Am J Physiol 273, C1378–1385.

Wu, S.K., Gomez, G.A., Michael, M., Verma, S., Cox, H.L., Lefevre, J.G., Parton, R.G., Hamilton, N.A., Neufeld, Z., and Yap, A.S. (2014). Cortical F-actin stabilization generates apical-lateral patterns of junctional contractility that integrate cells into epithelia. Nat Cell Biol 16, 167–178.

Yang, G., Truong, L.D., Timme, T.L., Ren, C., Wheeler, T.M., Park, S.H., Nasu, Y., Bangma, C.H., Kattan, M.W., Scardino, P.T., et al. (1998). Elevated expression of caveolin is associated with prostate and breast cancer. Clin Cancer Res 4, 1873–1880.

Yeow, I., Howard, G., Chadwick, J., Mendoza-Topaz, C., Hansen, C.G., Nichols, B.J., and Shvets, E. (2017). EHD Proteins Cooperate to Generate Caveolar Clusters and to Maintain Caveolae during Repeated Mechanical Stress. Curr Biol 27, 2951–2962 e2955.

