## Supplementary Materials for "Caveolae set levels of epithelial monolayer tension to eliminate tumor cells"

### SUPPLEMENTAL MATERIAL

#### Materials and Methods

*Cell culture, Transfection and Plasmids*

Mouse liver epithelial cells, AML12, were cultured in DMEM:F12 GlutaMAX™ medium supplemented with 10% FBS, 1% NEAA, 10μg/ml insulin, 5.5μg/ml transferrin, 5ng/ml selenium, 40ng/ml dexamethasone, 100 units/ml Penicillin and 100 units/ml Streptomycin. Human Embryonic Kidney 293 cells, HEK-293T, were cultured in DMEM medium supplemented with 10% FBS, 1% L-glutamine, 100 units/ml Penicillin and 100 units/ml Streptomycin. AML12 cells were transfected using Lipofectamine 3000 according to manufacturer's recommendation. HEK-293T cells were transfected using Lipofectamine 2000 (Invitrogen) for lentiviral expression vector pLL5.0 and third generation packaging constructs pMDLg/pRRE, RSV-Rev and pMD.G. Third generation packaging constructs were kindly provided by Prof. Jim Bear (UNC Chapel Hill, North Carolina, USA). To generate Cherry-HRas$^{V12}$ stable cells, we transduced AML12 cells with puromycin-resistant Cherry-HRas$^{V12}$ pTripZ lentiviral plasmid encoding the rtTA protein. The transduced AML12 cells were selected in media containing 3μg/ml of puromycin before inducing HRas$^{V12}$ expression with 2μg/ml of doxycycline.

Lentivirus-based shRNA was used to downregulate Caveolin-1 (CAV1), Cavin-1, Fmnl2, Fmnl3 and Inf2 in AML12 cells. shRNA sequences containing a stem loop sequence targeting the 3' UTR was designed and cloned into pLL5.0 vectors encoding fluorescent proteins such as EGFP, tagRFPT and NLSCherry whose expression is driven by a LTR promoter. shRNA sequences were cloned downstream of the U6 promoter using *HpaI* and *XhoI* giving rise to pLL5.0 ShCAV1-EGFP (5'-GCATCTTTCTCTCTGTTAA-3'), pLL5.0 ShCavin1-EGFP (5'-TGAGCAGCCGGACCAATAA-3'), pLL5.0 ShFmnl2-EGFP (5'-TGAGAGGTCTTCTAAGAAA-3'), pLL5.0 ShFmnl3-EGFP (5'-AGAAGGCTGTGTTGTTCAA-3') and pLL5.0 ShInf2-EGFP (5'-TGAGGGAATGCTCGTTGGA-3'). *Canis Lupus* CAV1 in pEGFP-N1, *Mus Musculus* Cavin-1 in pEGFP-C1, KRasG12V in pEGFP-N1 and c-Src$^{Y527F}$ in pEGFP-N1 plasmid were used as previously described (Ariotti et al., 2014; Hill et al., 2008; Liang et al., 2017). PLCδ-PH EGFP plasmid was kindly provided by Dr Markus Kerr (University of Queensland, Australia). Tet-on Cherry-HRas$^{V12}$ pTripZ lentiviral plasmid encoding the rtTA protein was generated by Dr Srikanth Budnar (Yap lab). Tet-on c-Src$^{Y527F}$ –Cherry pTripZ lentiviral plasmid was generated by synthesizing c-Src$^{Y527F}$ –Cherry flanked with AgeI and ClaI restriction sites before cloning the synthesized gene into pTripZ lentiviral plasmid digested with AgeI and ClaI. Human Fmnl2 in pEGFP-N1 was kindly provided by Prof Klemens Rottner (TU Braunschweig, Germany). pLL5.0 shCAV1-EGFP rescue construct was generated by cloning full length *Canis Lupus* CAV1 downstream of the LTR promoter before EGFP. This was similarly done for pLL5.0 shCavin1-EGFP rescue construct and pLL5.0 shFmnl2-EGFP rescue construct. pLL5.0 shEcad-EGFP rescue construct was also generated for laser ablation assays as mentioned above using the 3'UTR targeting sequence (5'-GGGAGATGCAGAATAATTA-3').

*Antibodies and Inhibitors*

Antibodies used in this project were: rabbit polyclonal antibody against CAV1 (N-20) (Santa Cruz, cat# SC894, 1:7000), rabbit polyclonal antibody against CAV1 (Abcam, cat# 2910, 1:200), rabbit polyclonal antibody against cavin1/PTRF (Sigma, cat# HPA049838, 1:2000), rabbit polyclonal antibody against cavin1 (C1, 1:500) (Bastiani et al., 2009) mouse monoclonal antibody against beta-tubulin (Sigma, cat# T4026, 1:2000), rat monoclonal antibody against E-cadherin (Invitrogen, cat# 131900, 1:400), rabbit polyclonal antibody against GAPDH (R&D systems, cat# 2275-PC-100, 1:10,000), mouse monoclonal antibody against GFP (Sigma, cat# 11814460001, 1:5000), rabbit polyclonal antibody against tagRFPT (Evrogen, cat# AB233, 1:2000), mouse monoclonal antibody against Fmnl2 (Abcam, cat# ab57963, 1:1000), rabbit polyclonal antibody against Fmnl2 (Sigma, cat# HPA005464, 1:1000), rabbit polyclonal antibody against Fmnl3 (Novus Biologicals, cat# NBP1-84092, 1:1000), rabbit polyclonal antibody against Inf2 (Proteintech, cat# 20466-1-AP, 1:1000), mouse monoclonal antibody against ZO-1 (Invitrogen, cat# 339100, 1:200), rabbit polyclonal antibody against ZO-1 (Invitrogen cat# 617300, 1:200), rabbit polyclonal against mCherry (Biovision, cat# 5993, 1:1000), rat monoclonal antibody against mCherry (Invitrogen, cat# M11217, 1:200), rabbit monoclonal antibody against Ras (Cell Signaling, cat# 3339 (27H5), 1:1000), rabbit polyclonal antibody against c-Src (Santa Cruz, cat# sc-18, 1:500), rabbit monoclonal antibody against myosin light chain 2 (Cell Signaling, cat# 8505 (D18E2), 1:1000), mouse monoclonal antibody against phospho-myosin light chain 2 (Ser19) (Cell Signaling, cat# 3675, 1:1000), mouse monoclonal antibody against p120 catenin (BD Transduction Laboratories, cat# 610134, 1:1000), rabbit polyclonal antibody against $\alpha$-catenin (Invitrogen, cat# 711200, 1:1000), mouse monoclonal antibody against β-catenin (BD Biosciences, cat# 610154, 1:2000), rabbit polyclonal antibody against myosin IIA (Sigma, cat# M8064, 1:1000), rabbit polyclonal antibody against myosin IIB (Sigma, cat# M7939, 1:1000), AlexaFluor 488-Phalloidin, AlexaFluor 546-Phalloidin (Invitrogen, 1:500) was used to stain for F-actin. Species-specific secondary antibodies conjugated with AlexaFluor 488, 546 or 647 (Invitrogen, 1:500) were used for immunofluorescence, or conjugated with horseradish peroxidase (BioRad Laboratories, 1:5000) for western blots. Cells were treated with 20μM SMIFH2 (TOCRIS, cat# 4401), 200nM Jasplakinolide (Merck, cat# 420107), 3mM Neomycin (Sigma, cat# N6386), 40μM RhoBQRL (custom synthesized, Biomatik USA) and PBP10 (TOCRIS, cat# 4611).

*Gene expression analysis of formin family members by quantitative PCR (QPCR)*

Total RNA from AML12 cells was isolated using GenElute™ Mammalian Total RNA Miniprep Kit (Sigma, RTN70) according to manufacturer's instructions. cDNA was synthesized from RNA using SuperScript™ III First-Strand Synthesis System (Life Technologies, cat# 18080051) according to manufacturer's instructions and cDNA was then used as a template for QPCR using SYBR™ Green PCR Master Mix (Life Technologies, cat# 4309155) and the Applied Biosystems ViiA 7 Real-Time PCR system (Life Technologies). Primers for housekeeping gene GAPDH and various formin members, as well as PCR cycling parameters were obtained and performed as published previously (Rosado et al., 2014). Relative quantification was performed using the comparative Ct method.

*Immunofluorescence microscopy*

Cells were fixed with 4% paraformaldehyde in cytoskeleton stabilization buffer (10 mM PIPES at pH 6.8, 100 mM KCl, 300 mM sucrose, 2 mM EGTA and 2 mM MgCl2) on ice for 30 min. PFA-fixed cells were permeabilized with 0.1% Saponin for 5 min on ice and subsequently blocked with 3% BSA for 1 hour prior to overnight primary antibody incubation. Subsequently, cells were washed with PBS and incubated with secondary antibodies for 1 hour before mounting with Prolong Gold Antifade Reagent with DAPI (Cell Signaling Technologies, cat# 8961). Cells were also fixed with ice-cold methanol for 5 min on ice.

Confocal images were acquired with an inverted Zeiss 880 or an upright 710 Meta laser-scanning microscope. Super resolution images were obtained with an inverted Structured Illumination microscope (SIM) with a PCO sCMOS camera at Queensland

Brain Institute (QBI) Advanced microimaging and analysis facility. Live-cell imaging was performed with an Nikon Ti-E Inverted microscope with Hamamatsu Flash 4.0 sCMOS camera. Alternatively, spinning disc confocal live cell imaging was performed on an environmentally controlled Nikon Ti inverted microscope with a Borealis-modified Yokogawa CSU-X1 confocal head (Spectral Applied Research) and equipped with an ASI PZ-2150 XYZ automated stage with piezo z-axis top plate, Clara cooled scientific grade interline CCD (Charge coupled device) camera (Andor) (Stehbens et al., 2012). SDC microscope hardware was controlled by NIS Elements software (Nikon). Subcellular distribution of E-cadherin, Caveolin-1 and ZO-1 was visualized by confocal fluorescence illumination on a Leica SP8 STED 3X FLIM Super Resolution instrument equipped with 405 nm, 440 nm pulsed, 442 nm and fully tunable supercontinuum white (470-670 nm) lasers, 592 nm, 660 nm and 775 nm STED lasers, PMT and high-sensitivity HyD detectors and controlled by Leica LASX software. Images were acquired using HC Plan Apochromat 100x1.40 oil objective and deconvolved using Huygens 17.04.1p1. Z-series were acquired using 0.1-mm-step confocal based scans, adjusted for brightness and presented as a 3D reconstruction using Imaris 9, or maximum projections using ImageJ 1.52g. Fluorescence intensity at cell junctions were quantified using line scan in ImageJ as described previously (Smutny et al., 2010).

*Quantification of apical cell extrusion and cell clusters*

Cells were mixed at a ratio of 1:20 and, to quantify the percentage of apical cell extrusion from the epithelial monolayer, we counted the number of cherry-positive cells that were 1) rounded up and above the plane of the monolayer 2) surrounded by cells that were not cherry-positive.

Cells were mixed at a ratio of 1:20 and 2ug/ml of doxycycline was added 24 hours later and the drug-containing medium refreshed daily. The number of cell clusters were counted at day 1 and day 4. Cell clusters were grouped into 4 categories: 1-2 cells, 3-4 cells, 5-10 cells and >10 cells. Quantification of number of >10 cells cell cluster was represented as a percentage between GFP control, ShCAV1 GFP and CAV1 Rescue GFP mixed with Tet-on Cherry-HRas$^{V12}$ –expressing WT cells. % Cell Cluster = [(number of clusters with > 10 cells) / (number of clusters with 1-2 cells + number of clusters with 3-4 cells + number of clusters with 5-10 cells + number of clusters with >10 cells)] *100.

*Laser Ablation*

AML12 cells are co-transduced with lentivirus encoding pLL5.0 ShCAV1 GFP or pLL5.0 KnockdownRescueCAV1GFP with pLL5.0 KnockdownRescueE-cadherin-TagRFPT. For live-cell experiments, cells were cultured on 29 mm glass-bottom dishes (Shengyou Biotechnology) and imaged in clear Hank's balanced salt solution supplemented with 5% FBS, 10 mM HEPES (pH 7.4) and 5 mM $CaCl_2$. Cells were processed for experiments 48 hours post transduction. Laser ablation experiment was performed on an LSM 510 meta Zeiss confocal microscope equipped with a 37°C heating stage. Images were acquired with a 63× objective (1.4 NA oil Plan Apochromat immersion lens) with 1.5× zoom. A constant ROI, 3.8 X 0.6μm, with the longer axis parallel to the cell–cell contact was ablated with a Ti:sapphire laser (Chameleon Ultra, Coherent Scientific) tuned to 790 nm, 30 iterations, and 22% transmission. Time-lapse images were acquired before (2 frames, 5.28s) and after (21 frames, 114.72s) ablation. Image analysis was done as described previously (Liang et al., 2016). Briefly, the distance between vertices of the cell-cell junctions was tracked in ImageJ and measured as a function of time. The mean values of tracked distances were plotted against time to obtain the initial recoil. At least 17 junctions were ablated and calculated for each condition in every experiment. The cell-cell junctions were modelled as a Kelvin-Voigt fiber (Fernandez-Gonzalez et al., 2009) and their deformation [L(t) – L (0)] was described by the following equation:

$$L(t) - L(0) = \frac{Fo}{E} \ .(1 - \mathrm{e}^{\frac{-E}{\mu}.t})$$

F refers to the force applied to the junction before ablation, E refers to the elastic modulus of the junction and μ refers to the viscosity coefficient related to the viscous drag of the media. Initial recoil velocity was calculated as:

$$Initial\ Recoil = \frac{d[L(t) - L(0)]}{dt} = \frac{Fo}{\mu}$$

Initial recoil reflects the tensile force applied at junctions prior to the ablation of the junctions, assuming a constant viscosity coefficient among experiments. To assess changes in junctional elasticity, we calculated the viscosity coefficient as:

$$k = \frac{E}{\mu}$$

k values that are not significantly altered suggest that the changes of initial velocity were attributed to the changes of tensile force.

*Fluorescence Recovery After Photobleaching (FRAP) and Photoactivation and FRET analysis*

FRAP and photoactivation experiments were performed on an inverted Zeiss Meta LSM 710 confocal microscope with an incubation box maintained at 37$^{o}$C. Time-lapse images were captured using a 63X objective before and after photobleaching with a 488nm laser, at an interval of approximately 7 seconds per frame. Image analysis was done as described previously (Priya and Gomez). For photoactivation experiments, protein was photoactivated with a Mai Tai laser with 2% transmission and 30 iterations. Time-lapse images were captured before and after photoactivation at an interval of 93 millisecond per frame. Image analysis was done as previously described (Wu et al., 2014). FRET measurement was performed as published previously (Acharya et al., 2017).

*Quantification of actin bundle organization*

To determine the number of intersects formed per area, SIM images of F-actin filaments were analysed using FIJI. A region of interest was drawn over junctional F-actin and a bounding box was generated. Following background subtraction, thresholding was performed to generate a binary image. Skeletonize 2D/3D plugin was applied to the image followed by the Analyze Skeleton plugin (Michael et al., 2016). For nematic order parameter analysis of F-actin organization, we used Matlab codes kindly provided by Prof Stephan W. Grill (Reymann et al., 2016). Briefly, a cell junction was randomly segmented in square templates of 2.64μm and subjected to Fourier transform analysis of F-actin bundles for quantification.

*Electron Microscopy Images*

Fixed cell monolayers were processed in dishes following the method by Tapia *et al.* (Tapia et al., 2012). Cells were incubated in 2% osmium tetroxide and 1.5% potassium ferricyanide (aqueoussolution) for 1 hour at room temperature, followed by a 1% aqueous thiocarbohydrazide solution for 20min. Cells underwent further osmification in 2% osmium tetroxide (aqueous) for 30min, again at room temperature. Cells were exchanged into 1% uranyl acetate overnight at 4°C. The following day the cells were placed in lead aspartate for 60 min at 60°C. Between each staining step the cells were washed thoroughly in UHQ water (3 x 5min).

Samples were dehydrated with increasing concentrations of ethanol (20%, 50%, 70%, 90%, 3 x 100%) for 5 min each and infiltrated with increasing concentrations of Durcupan resin in ethanol for 2 hours each (25%, 50%, and 75%). Samples were left in 100% Durcupan resin overnight and placed in fresh resin for 2 hours the following day. The majority of the resin was removed from the dish until a fine layer remained, the dish was them placed in the oven at 60°C for 48 hours to polymerise. The resin/cell layer was removed from the dish and a small piece, where cell growth was highly confluent, was mounted onto SEM stubs using two-part silver expoy.

Data sets were analysed using Imod software (Kremer et al., 1996). Image stacks were aligned manually using the Midas command. Caveolae were identified and marked with a single point using the manual drawing tool. The use of the scattered points options allowed a spherical mesh to be created for each point.

*Traction Force microscopy and Intercellular Stress measurement*

Traction force microscopy was performed as previously described (Saw et al., 2017). Briefly, soft silicone gel with a stiffness of 10-20kPa was prepared by mixing CyA and CyB at 1:1 ratio (Dow Corning). The substrate was cured at 80°C for 2 hours before silanizing with 5% 3-aminopropyl trimethoxysilane (Sigma) in ethanol. 0.05% of 200nm carboxylated fluorescent beads were added to the substrate and the substrate was coated with 10μg/ml of fibronectin. Subsequently, cells were seeded on the substrate and live-cell imaging was performed. Cells were imaged with a 20X objective, capturing an image every 7 minutes for 12 hours. After 12 hours, 10% SDS was added to remove cells and bead images were captured during the resting state of the gel. Bead displacement measurements were converted to traction forces using an ImageJ plugin (Martiel et al., 2015) and intercellular stress measurements were performed using the BISM MATLAB script (Nier et al., 2016).

**Supplementary Figure Captions**

**Figure S1 related to Fig 1.**

(a) Immunofluorescence and western blot showing endogenous CAV1 and CAV1 GFP fusion proteins in GFP control, ShCAV1 GFP and CAV1 Rescue GFP cells.

(b) Electron micrographs of AML12 cells with caveolae highlighted by red markers in the right panel (unmarked image in left panel).

(c) Deconvolved confocal images of AML12 cells co-stained with E-cadherin, ZO-1 and CAV1 (Scale bar = 4μm).

(d) FIB-SEM images of AML12 cells, with caveolae highlighted by blue markers.

(e) Immunofluorescence and quantification of endogenous junctional Cavin-1 staining in GFP control, ShCAV1 GFP and CAV1 Rescue GFP cells.

(f) Immunofluorescence and western blot showing endogenous levels of Cavin1 and Cavin1 GFP fusion proteins in GFP control, ShCavin1 GFP and Cavin1 Rescue GFP cells.

(g) Quantification of apical oncogenic cell extrusion of single cells expressing $KRas^{G12V}$ within GFP control, ShCAV1 GFP and CAV1 Rescue GFP epithelial monolayers.

(h) Schematic illustration and quantification of single $c\text{-}Src^{Y527F}$-expressing cells within GFP control, ShCavin1 GFP and Cavin1 Rescue GFP epithelial monolayers.

(i) Western blot of endogenous CAV1 levels in GFP control, ShCavin1 GFP and Cavin1 Rescue GFP cells. Western blot of endogenous Cavin1 levels in wild-type, GFP control, ShCAV1 GFP and CAV1 Rescue GFP cells.

All data are means ± s.e.m; n.s, not significant; **P*<0.05, ***P*<0.01, ****P*<0.001, *****P*<0.0001; calculated from N=3 independent experiments analysed with two-way ANOVA Tukey's multiple comparisons test except (d) which was analysed with one-way ANOVA, Tukey's multiple comparisons test. Scale bars, 10μm.

**Figure S2 related to Figure 3.**

(a) Junctional recoil curve of GFP control, ShCavin1 GFP and Cavin1 Rescue GFP following laser ablation.

(b) Western blots of $Tet^{on}$ Cherry-$HRas^{V12}$ probled for Ras or the mCherry epitope tag before and 24hrs post doxycycline treatment.

(c) Overexpression of MRLC-EGFP in AML12 cells.

(d) Western blot of total MLC, phosphoMLC and GFP in GFP control and GFP-MRLC cells.

(e) Quantification of pMLC/Total MLC in GFP control and GFP-MRLC cells.

(f) Recoil curve and (g) quantification of initial recoil of GFP control and GFP-MRLC cells following laser ablation.

(h) Representative images and quantification of junctional Cavin1 in GFP control and GFP-MRLC cells.

(i) Western blot of $Tet^{on}$ c-$Src^{Y527F}$-Cherry before and 24hrs post doxycycline treatment.

(j) Quantification of apical oncogenic cell extrusion in mixing experiments with $Tet^{on}$ c-$Src^{Y527F}$-expressing WT cells surrounded by GFP control or GFP-MRLC cells.

All data are means ± s.e.m; n.s, not significant; **P*<0.05, ***P*<0.01, ****P*<0.001, *****P*<0.0001; calculated from N=3 independent experiments analysed with Student's *t*-test. Scale bar, 10μm.

**Figure S3 related to Figure 4.**

(a-c) Quantification of junctional myosin IIA (a), (b) myosin IIB and (c) E-cadherin in GFP control, ShCAV1 GFP and CAV1 Rescue GFP cells.

(d) Western blot of endogenous E-cadherin and catenins in GFP control, ShCAV1 GFP and CAV1 Rescue GFP cells.

(e) Fluorescence recovery curve of E-cadherin-EGFP in GFP control, ShCAV1 GFP and CAV1 Rescue GFP cells.

(f,g) Quantification of Half-life (f) and (g) mobile fraction of E-cadherin-EGFP calculated from FRAP analysis of GFP control, ShCAV1 GFP and CAV1 Rescue GFP cells.

All data are means ± s.e.m; n.s, not significant; **P*<0.05, ***P*<0.01, ****P*<0.001, *****P*<0.0001; calculated from N=3 independent experiments analysed with one-way ANOVA Dunnett's multiple comparisons test except (f) and (g) which were analysed with one-way ANOVA, Tukey's multiple comparisons test.

**Figure S4 related to Figure 5.**

(a) mRNA expression level of formins in GFP control cells. n.d., not determined.

(b) Relative expression of formin mRNA in GFP control, ShCAV1 GFP and CAV1 Rescue GFP cells measured by QPCR.

(c) Western blot of endogenous FMNL2/3 in GFP control, ShCAV1 GFP and CAV1 Rescue GFP cells.

(d) Western blot of endogenous FMNL2, Fmnl3, INF2 and CAV1 in control, ShCAV1, ShCav1 downregulated for FMNL2 or INF2 or FMNL3.

(e) Junctional recoil curves of control, ShCav1 and ShCav1 downregulated for FMNL2 following laser ablation.

**Figure S5 related to Figure 6.**

(a) Representative images of tagRFPT control, ShCAV1 tagRFPT and CAV1 Rescue tagRFPT cells transfected with YFP-CRIB location biosensor to detect endogenous Cdc42-GTP.

(b) Quantification of normalized junctional intensity of PLC$\delta$-PH in control, ShCAV1 and CAV1 Rescue cells treated with and without Neomycin.

(c) PBP10 masks access to PtdIns(4,5)$P_2$. Quantification of junctional intensity of PLC$\delta$-PH in control, ShCAV1 and CAV1 Rescue cells treated with and without PBP10.

(d) Quantification of junctional intensity of PLC$\delta$-PH in control, ShCAV1 and CAV1 Rescue cells treated with and without RhoB-QRL.

(e) Representative images and (f) quantification of normalized junctional intensity of FMNL2-EGFP in control, ShCAV1 and CAV1 Rescue cells treated with and without PBP10.

(g) Quantification of junctional intensity of FMNL2-EGFP in control, ShCAV1 and CAV1 Rescue cells treated with and without RhoB-QRL.

(h) Representative images of junctional FMNL2$^{G2A}$, FMNL2$^{12A}$ and FMNL2$^{G2A\text{-}12A}$.

(i) Western blot of endogenous CAV1, FMNL2 and FMNL2-EGFP mutant construct.

(j) Recoil curve of ShCAV1 co-transduced by GFP control, ShFMNL2, FMNL2$^{WT}$ and FMNL2$^{12A}$ derived from laser ablation. All data are means $\pm$ s.e.m; n.s, not significant; **P*<0.05, ***P*<0.01, ****P*<0.001, *****P*<0.0001; calculated from N=3 independent experiments analysed with two-way ANOVA Tukey's multiple comparisons test except (f) which was analysed with Sidak's multiple comparisons test. Scale bar, 10$\mu$m.

**Figure S6 related to Figure 7.**

(a) Quantification of apical oncogenic cell extrusion of HRas$^{V12}$-transfected control and ShCAV1 epithelial monolayers treated with and without SMIFH2.

(b) Stacked column graph indicating the number of $Tet^{on}$ $HRas^{V12}$ cell clusters in control, ShCAV1 and CAV1 Rescue epithelial; 1 day and 4 days after doxycycline induction.

All data are means ± s.e.m; n.s, not significant; **P*<0.05, ***P*<0.01, ****P*<0.001, *****P*<0.0001; calculated from N=3 independent experiments analysed with two-way ANOVA Tukey's multiple comparisons test**.**

**Supplementary Table 1 related to Figure 3, 4, 5, 6.**

k values obtained from junctional recoil measurements from laser ablation. All data are means ± SEM; ns, not significant; **P*<0.05, ***P*<0.01, ****P*<0.001, *****P*<0.0001; calculated from N=3 independent experiments. Data analysed with one-way ANOVA Tukey's multiple comparison's test except for SMIFH2 treatment which were analysed with two-way ANOVA Tukey's multiple comparison's test, and GFP control and MRLC-EGFP which were analysed with Student's *t*-test.

**References**

Acharya, B.R., Wu, S.K., Lieu, Z.Z., Parton, R.G., Grill, S.W., Bershadsky, A.D., Gomez, G.A., and Yap, A.S. (2017). Mammalian Diaphanous 1 Mediates a Pathway for E-cadherin to Stabilize Epithelial Barriers through Junctional Contractility. Cell Rep *18*, 2854-2867.

Ariotti, N., Fernandez-Rojo, M.A., Zhou, Y., Hill, M.M., Rodkey, T.L., Inder, K.L., Tanner, L.B., Wenk, M.R., Hancock, J.F., and Parton, R.G. (2014). Caveolae regulate the nanoscale organization of the plasma membrane to remotely control Ras signaling. J Cell Biol *204*, 777-792.

Bastiani, M., Liu, L., Hill, M.M., Jedrychowski, M.P., Nixon, S.J., Lo, H.P., Abankwa, D., Luetterforst, R., Fernandez-Rojo, M., Breen, M.R.*, et al.* (2009). MURC/Cavin-4 and cavin family members form tissue-specific caveolar complexes. J Cell Biol *185*, 1259-1273.

Fernandez-Gonzalez, R., Simoes Sde, M., Roper, J.C., Eaton, S., and Zallen, J.A. (2009). Myosin II dynamics are regulated by tension in intercalating cells. Dev Cell *17*, 736-743.

Hill, M.M., Bastiani, M., Luetterforst, R., Kirkham, M., Kirkham, A., Nixon, S.J., Walser, P., Abankwa, D., Oorschot, V.M., Martin, S.*, et al.* (2008). PTRF-Cavin, a conserved cytoplasmic protein required for caveola formation and function. Cell *132*, 113-124.

Kremer, J.R., Mastronarde, D.N., and McIntosh, J.R. (1996). Computer visualization of three-dimensional image data using IMOD. J Struct Biol *116*, 71-76.

Liang, X., Budnar, S., Gupta, S., Verma, S., Han, S.P., Hill, M.M., Daly, R.J., Parton, R.G., Hamilton, N.A., Gomez, G.A.*, et al.* (2017). Tyrosine dephosphorylated cortactin downregulates contractility at the epithelial zonula adherens through SRGAP1. Nat Commun *8*, 790.

Liang, X., Michael, M., and Gomez, G.A. (2016). Measurement of Mechanical Tension at Cell-cell Junctions Using Two-photon Laser Ablation. Bio Protoc *6*.

Martiel, J.L., Leal, A., Kurzawa, L., Balland, M., Wang, I., Vignaud, T., Tseng, Q., and Thery, M. (2015). Measurement of cell traction forces with ImageJ. Methods Cell Biol *125*, 269-287.

Michael, M., Meiring, J.C.M., Acharya, B.R., Matthews, D.R., Verma, S., Han, S.P., Hill, M.M., Parton, R.G., Gomez, G.A., and Yap, A.S. (2016). Coronin 1B Reorganizes the Architecture of F-Actin Networks for Contractility at Steady-State and Apoptotic Adherens Junctions. Dev Cell *37*, 58-71.

Nier, V., Jain, S., Lim, C.T., Ishihara, S., Ladoux, B., and Marcq, P. (2016). Inference of Internal Stress in a Cell Monolayer. Biophys J *110*, 1625-1635.

Priya, R., and Gomez, G.A. (2013). Measurement of Junctional Protein Dynamics Using Fluorescence Recovery After Photobleaching (FRAP). Bio-protocol *3*, e937.

Reymann, A.C., Staniscia, F., Erzberger, A., Salbreux, G., and Grill, S.W. (2016). Cortical flow aligns actin filaments to form a furrow. Elife *5*.

Rosado, M., Barber, C.F., Berciu, C., Feldman, S., Birren, S.J., Nicastro, D., and Goode, B.L. (2014). Critical roles for multiple formins during cardiac myofibril development and repair. Mol Biol Cell *25*, 811-827.

Saw, T.B., Doostmohammadi, A., Nier, V., Kocgozlu, L., Thampi, S., Toyama, Y., Marcq, P., Lim, C.T., Yeomans, J.M., and Ladoux, B. (2017). Topological defects in epithelia govern cell death and extrusion. Nature *544*, 212-216.

Smutny, M., Cox, H.L., Leerberg, J.M., Kovacs, E.M., Conti, M.A., Ferguson, C., Hamilton, N.A., Parton, R.G., Adelstein, R.S., and Yap, A.S. (2010). Myosin II isoforms identify distinct functional modules that support integrity of the epithelial zonula adherens. Nat Cell Biol *12*, 696-702.

Stehbens, S., Pemble, H., Murrow, L., and Wittmann, T. (2012). Imaging intracellular protein dynamics by spinning disk confocal microscopy. Methods Enzymol *504*, 293-313.

Tapia, J.C., Kasthuri, N., Hayworth, K.J., Schalek, R., Lichtman, J.W., Smith, S.J., and Buchanan, J. (2012). High-contrast en bloc staining of neuronal tissue for field emission scanning electron microscopy. Nat Protoc *7*, 193-206.

Wu, S.K., Gomez, G.A., Michael, M., Verma, S., Cox, H.L., Lefevre, J.G., Parton, R.G., Hamilton, N.A., Neufeld, Z., and Yap, A.S. (2014). Cortical F-actin stabilization generates apical-lateral patterns of junctional contractility that integrate cells into epithelia. Nat Cell Biol *16*, 167-178.

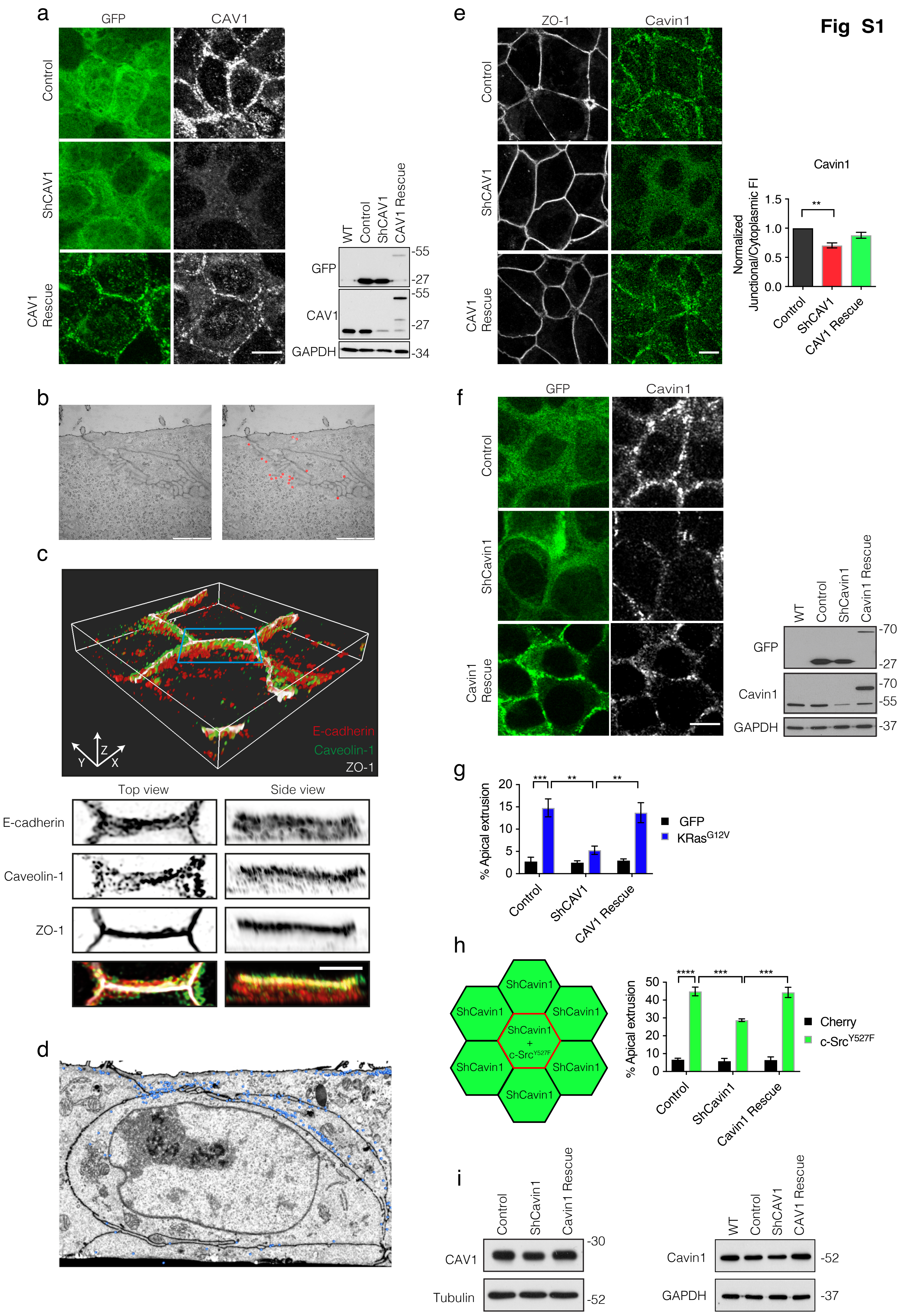
Fig S1
a
GFP
CAV1
Control
ShCAV1
CAV1 Rescue
WT
Control
ShCAV1
CAV1 Rescue
GFP
CAV1
GAPDH
-55
-27
-55
-27
-34
b
c
E-cadherin
Caveolin-1
ZO-1
Z
Y
X
Top view
Side view
E-cadherin
Caveolin-1
ZO-1
d
e
ZO-1
Cavin1
Control
ShCAV1
CAV1 Rescue
Cavin1
Normalized Junctional/Cytoplasmic FI
1.5
1.0
0.5
0.0
**
Control
ShCAV1
CAV1 Rescue
f
GFP
Cavin1
Control
ShCavin1
Cavin1 Rescue
WT
Control
ShCavin1
Cavin1 Rescue
GFP
Cavin1
GAPDH
-70
-27
-70
-55
-37
g
% Apical extrusion
20
15
10
5
0
***
**
**
GFP
KRas G12V
Control
ShCAV1
CAV1 Rescue
h
ShCavin1
ShCavin1 + c-Src Y527F
% Apical extrusion
50
40
30
20
10
0
****
***
***
Cherry
c-Src Y527F
Control
ShCavin1
Cavin1 Rescue
i
Control
ShCavin1
Cavin1 Rescue
CAV1
Tubulin
-30
-52
WT
Control
ShCAV1
CAV1 Rescue
Cavin1
GAPDH
-52
-37

### Fig S2

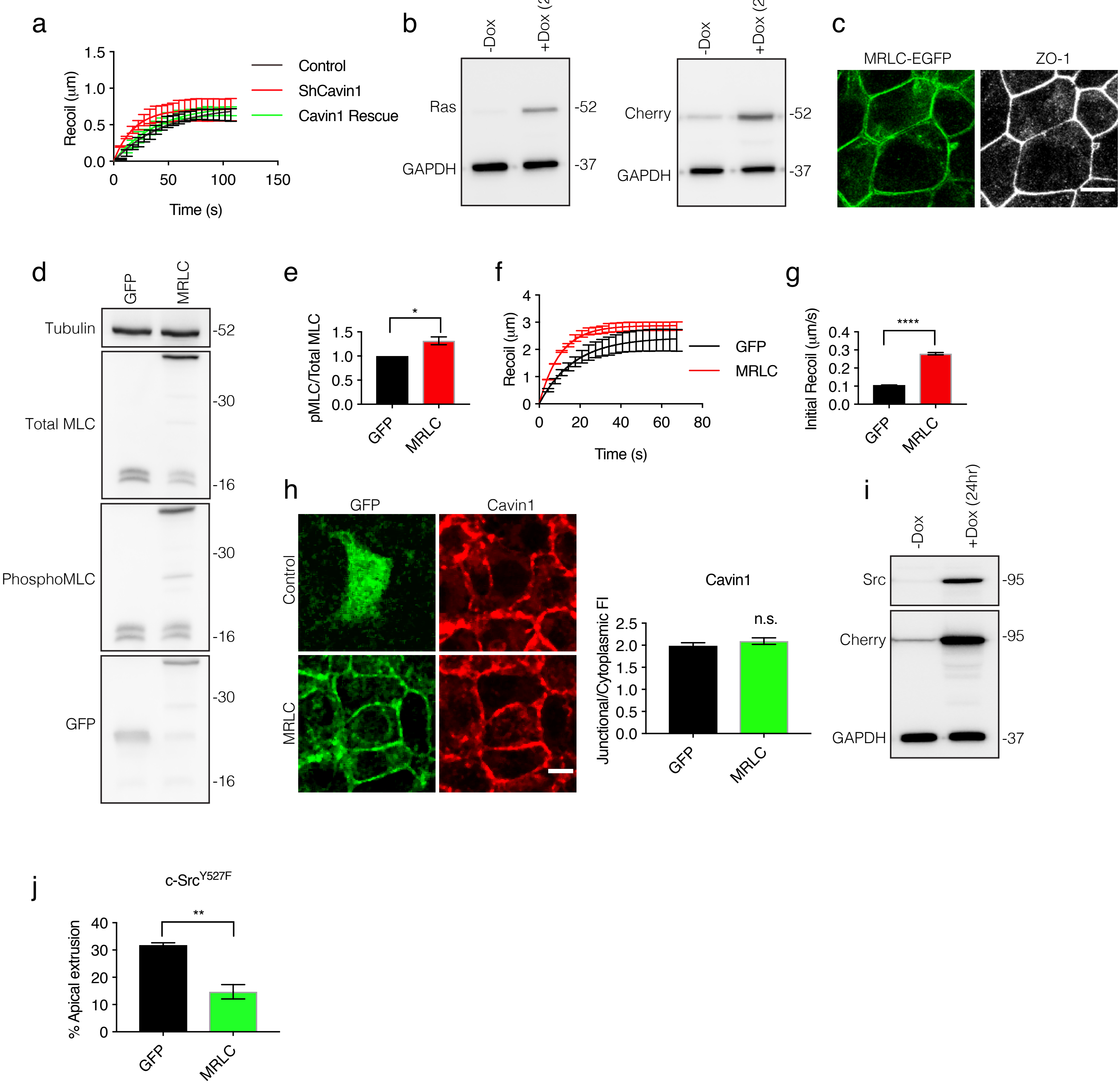
a
Recoil (µm)
Time (s)
Control
ShCavin1
Cavin1 Rescue
b
-Dox
+Dox (24hr)
Ras
GAPDH
Cherry
-52
-37
c
MRLC-EGFP
ZO-1
d
GFP
MRLC
Tubulin
Total MLC
PhosphoMLC
-52
-30
-16
e
pMLC/Total MLC
*
f
Recoil (µm)
Time (s)
GFP
MRLC
g
Initial Recoil (µm/s)
****
h
GFP
Cavin1
Control
MRLC
Junctional/Cytoplasmic FI
n.s.
i
-Dox
+Dox (24hr)
Src
Cherry
GAPDH
-95
-37
j
c-Src Y527F
% Apical extrusion
**

Fig S3

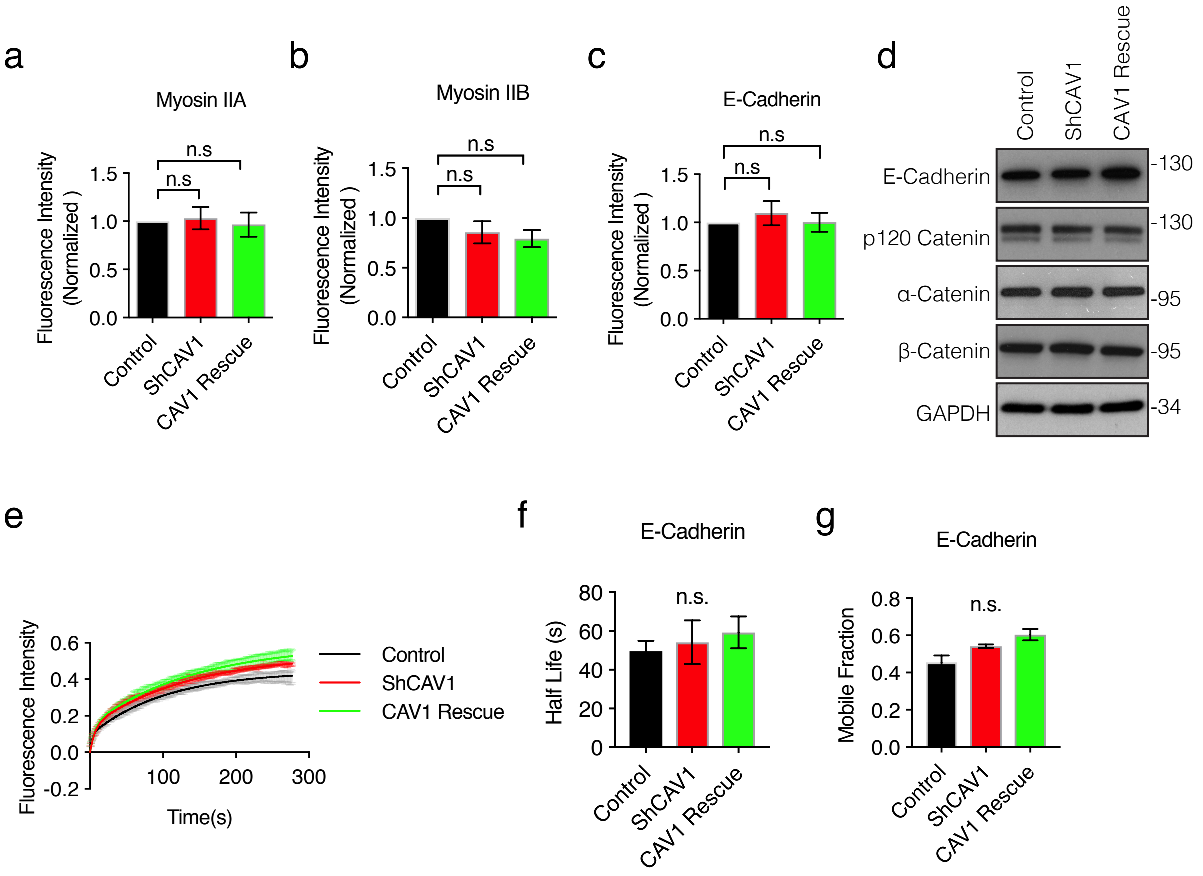
a
Myosin IIA
Fluorescence Intensity (Normalized )
n.s
n.s
1.5
1.0
0.5
0.0
Control
ShCAV1
CAV1 Rescue
b
Myosin IIB
Fluorescence Intensity (Normalized )
n.s
n.s
Control
ShCAV1
CAV1 Rescue
c
E-Cadherin
Fluorescence Intensity (Normalized )
n.s
n.s
Control
ShCAV1
CAV1 Rescue
d
Control
ShCAV1
CAV1 Rescue
E-Cadherin -130
p120 Catenin -130
α-Catenin -95
β-Catenin -95
GAPDH -34
e
Fluorescence Intensity
0.6
0.4
0.2
0.0
-0.2
100
200
300
Time(s)
Control
ShCAV1
CAV1 Rescue
f
E-Cadherin
Half Life (s)
80
60
40
20
0
n.s.
Control
ShCAV1
CAV1 Rescue
g
E-Cadherin
Mobile Fraction
0.8
0.6
0.4
0.2
0.0
n.s.
Control
ShCAV1
CAV1 Rescue

**Fig S4**

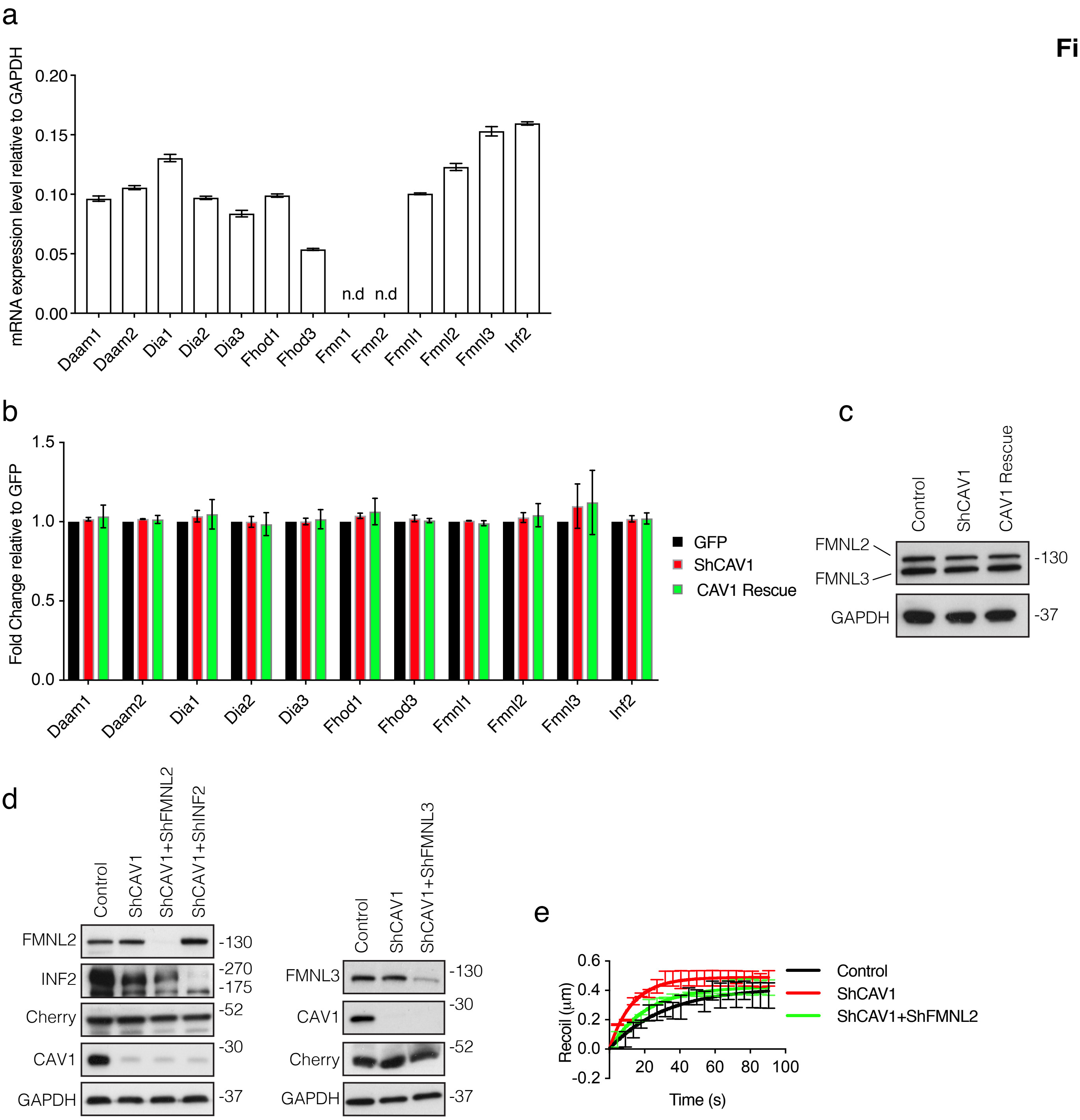
a
mRNA expression level relative to GAPDH
0.20
0.15
0.10
0.05
0.00
n.d
n.d
Daam1
Daam2
Dia1
Dia2
Dia3
Fhod1
Fhod3
Fmn1
Fmn2
Fmnl1
Fmnl2
Fmnl3
Inf2
b
Fold Change relative to GFP
1.5
1.0
0.5
0.0
GFP
ShCAV1
CAV1 Rescue
Daam1
Daam2
Dia1
Dia2
Dia3
Fhod1
Fhod3
Fmnl1
Fmnl2
Fmnl3
Inf2
c
Control
ShCAV1
CAV1 Rescue
FMNL2
FMNL3
-130
GAPDH
-37
d
Control
ShCAV1
ShCAV1+ShFMNL2
ShCAV1+ShINF2
FMNL2
-130
INF2
-270
-175
Cherry
-52
CAV1
-30
GAPDH
-37
Control
ShCAV1
ShCAV1+ShFMNL3
FMNL3
-130
CAV1
-30
Cherry
-52
GAPDH
-37
e
Recoil (μm)
0.6
0.4
0.2
0.0
-0.2
20
40
60
80
100
Time (s)
Control
ShCAV1
ShCAV1+ShFMNL2

**Fig S5**

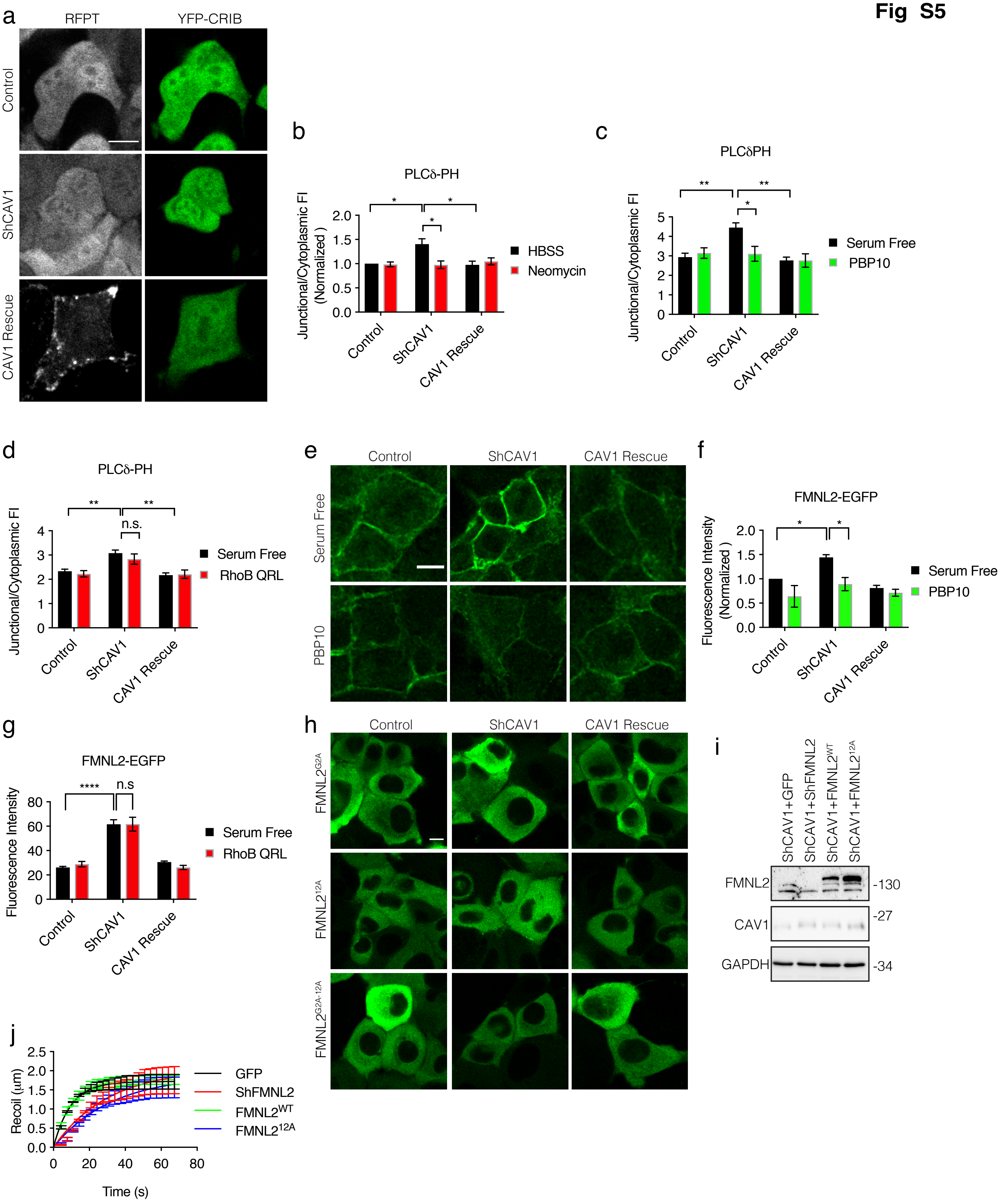
a
RFPT
YFP-CRIB
Control
ShCAV1
CAV1 Rescue
b
PLCδ-PH
Junctional/Cytoplasmic FI (Normalized )
HBSS
Neomycin
Control
ShCAV1
CAV1 Rescue
c
PLCδPH
Junctional/Cytoplasmic FI
Serum Free
PBP10
d
PLCδ-PH
n.s.
Junctional/Cytoplasmic FI
Serum Free
RhoB QRL
e
Control
ShCAV1
CAV1 Rescue
Serum Free
PBP10
f
FMNL2-EGFP
Fluorescence Intensity (Normalized )
Serum Free
PBP10
g
FMNL2-EGFP
n.s
Fluorescence Intensity
Serum Free
RhoB QRL
h
Control
ShCAV1
CAV1 Rescue
FMNL2G2A
FMNL212A
FMNL2G2A-12A
i
ShCAV1+GFP
ShCAV1+ShFMNL2
ShCAV1+FMNL2WT
ShCAV1+FMNL212A
FMNL2
CAV1
GAPDH
-130
-27
-34
j
Recoil (μm)
Time (s)
GFP
ShFMNL2
FMNL2WT
FMNL212A

**Fig S6**

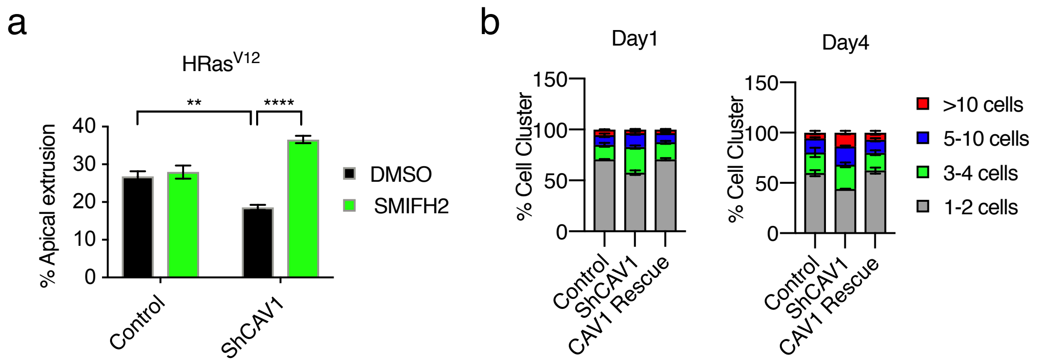
a
HRas$^{V12}$
**
****
% Apical extrusion
40
30
20
10
0
Control
ShCAV1
DMSO
SMIFH2
b
Day1
Day4
% Cell Cluster
150
100
50
0
Control
ShCAV1
CAV1 Rescue
>10 cells
5-10 cells
3-4 cells
1-2 cells

#### Table S1

| Condition | K Values | $p<0.05$ |
|---|---|---|
| pLL5.0 GFP | 0.0200 ± 0.01, N=3 | n.s |
| ShCAV1 GFP | 0.0339 ± 0.01, N=3 | |
| ResCAV1 GFP | 0.0252 ± 0.007, N=3 | |
| pLL5.0 GFP | 0.0217 ± 0.004, N=3 | n.s |
| ShCavin1 GFP | 0.0629 ± 0.01, N=3 | |
| ResCavin1 GFP | 0.0252 ± 0.005, N=3 | |
| pLL5.0 GFP_DMSO | 0.0938 ± 0.04, N=3 | n.s. |
| ShCAV1 GFP_DMSO | 0.1779 ± 0.07, N=3 | |
| pLL5.0 GFP_SMIFH2 | 0.0836 ± 0.01, N=3 | |
| ShCAV1 GFP_SMIFH2 | 0.0559 ± 0.02, N=3 | |
| pLL5.0 GFP | 0.0664 ± 0.02, N=3 | n.s |
| pLL5.0 MRLC-EGFP | 0.0996 ± 0.006, N=3 | |
| pLL5.0 NLSCherry_GFP | 0.0337± 0.009, N=3 | n.s |
| pLL5.0 ShCAV1NLSCherry_GFP | 0.0826 ± 0.01, N=3 | |
| pLL5.0 ShCAV1NLSCherry_ShFMNL2 GFP | 0.0528 ± 0.01, N=3 | |
| ShCAV1_pLL5.0 GFP | 0.1138 ± 0.02, N=3 | n.s. |
| ShCAV1_pLL5.0 ShFMNL2 | 0.0427 ± 0.01, N=3 | |
| ShCAV1_pLL5.0 FMNL2-WT | 0.1007 ± 0.008, N=3 | |
| ShCAV1_pLL5.0 ShFMNL2-12A | 0.0392 ± 0.01, N=3 | |
